# Structural analysis shows that the BIR2 domain of E3 ligase XIAP binds across the RIP2 kinase dimer interface

**DOI:** 10.1101/2022.10.14.512215

**Authors:** Mathilde Lethier, Michael Hons, Adrien Favier, Bernhard Brutscher, Elisabetta Boeri Erba, Stephen Cusack, Erika Pellegrini

## Abstract

NOD1 and NOD2 are innate immune system pattern recognition receptors that play a key role in the protection from bacterial infections and in the maintenance of gastro-intestinal homeostasis. Dysregulation of NOD signalling pathways promotes chronic inflammatory diseases such as inflammatory bowel disease. RIP2, which contains a kinase domain (RIP2K) and a CARD domain, is the immediate downstream signalling partner in the NOD pathway and inhibition of its kinase activity and ubiquitination are promising strategies to address these inflammatory diseases. However, recent work indicates that the phosphorylation activity of RIP2K is dispensable for signalling and that inhibitors of both RIP2K activity and RIP2 ubiquitination prevent the essential interaction between RIP2K and the BIR2 domain of XIAP, the key RIP2 ubiquitin E3 ligase. Moreover, XIAP BIR2 antagonists also block this interaction. To reveal the molecular mechanisms involved, we combined Native mass spectrometry, NMR, cryo-electron microscopy and Alphafold2 predictions to determine the structure of the RIP2K-XIAP BIR2 complex with 2:1 stoichiometry. The structure shows that complex formation requires that the kinase is in an active-like (i.e. αC-helix IN) and dimeric conformation and explains both inhibitory mechanisms. It also shows why phosphorylation of the kinase activation loop is dispensable for signalling and reveals the structural role of RIP2K-K209 residue in the RIP2K-XIAP BIR2 interaction. Importantly, our results clarify the features of the RIP2K conformation essential for its role as a scaffold protein for ubiquitination.

## Introduction

The innate immune system is a frontline defence strategy that allows vertebrates to combat infectious agents. Pattern Recognition Receptors (PRRs) that recognise non-self, molecular patterns and their signalling pathways are key elements of the innate immune system. Nucleotide-binding oligomerization domain-containing 1 and 2 (NOD1 and NOD2) proteins are intracellular PRRs sensing bacterial infections and contributing to gastro-intestinal homeostasis (Philpott *et al,* 2014; Trindade & Chen, 2020; Kayama Hisako, 2020). Receptor activation by bacterial peptidoglycan breakdown products, γ-D-Glu-mdiaminopimelic acid (iE-DAP) or muramyl-dipeptide (MDP) respectively (Girardin *et al,* 2016, 2003; Inohara *et al,* 2003; Chamaillard *et al,* 2003), triggers production of nuclear factor kappa B (NF-κB) and mitogen-activated protein kinase (MAPK) cascade activation. This results in transcriptional upregulation and release of pro-inflammatory cytokines and anti-microbial peptides (Boyle *et al,* 2014; Trindade & Chen, 2020; Philpott *et al,* 2014; Ferrand & Ferrero, 2013). Alternatively, NOD signalling can be triggered by endoplasmic reticulum (ER) stress and small Rho GTPase (Keestra & Bäumler, 2014; Keestra-Gounder *et al,* 2016). NOD activation also triggers autophagy, through the recruitment of autophagy related 16 like 1 (ATG16L1) (Travassos *et al,* 2010).

Dysregulated NOD signalling, which results in increased production of pro-inflammatory cytokines, impaired autophagy and therefore chronic inflammation is associated with several complex multifactorial inflammatory diseases, such as inflammatory bowel disease (IBDs), Blau syndrome and early-onset sarcoidosis (EOS). IBDs, which include Crohn’s disease (CD) and ulcerative colitis (UC), are a major social health problem in need of novel treatments (Kaplan & Windsor, 2021). Loss of function by mutations of NOD2 or ATG16L1 and misregulation of ubiquitination/deubiquitination in the NOD pathway can result in a genetic susceptibility to develop IBDs (Zou *et al,* 2021; Ananthakrishnan AN, 2015; Cho & Abraham, 2007; Philpott *et al,* 2014; Caruso, R., Lo, B.C. & Núñez, 2020; Kuballa *et al,* 2008; Cooney *et al,* 2010; Honjo *et al,* 2021a; Sámano-Sánchez & Gibson, 2020; Inohara *et al,* 2003). Dysregulation of NOD signalling has also been associated with other inflammatory disorders, e.g. inflammatory arthritis, asthma, colorectal cancer, multiple sclerosis and type 2 diabetes mellitus (Philpott *et al,* 2014; Caruso *et al,* 2014; Amar *et al,* 2011). For all these diseases, inhibition of NF-κB activation and therefore of pro-inflammatory cytokine production, has been suggested as a possible treatment (Boyle *et al,* 2014; Philpott *et al,* 2014; Trindade & Chen, 2020; Canning *et al,* 2015; Atreya *et al,* 2008).

Activation of the NOD receptors leads to the recruitment of receptor-interacting serine/threonine/tyrosine-protein kinase 2 (RIP2) (Boyle *et al,* 2014; Park *et al,* 2007). RIP2 is an adaptor protein, which comprises a kinase (RIP2K) and a Caspase activation and recruitment domain (CARD) connected by a disordered intermediate loop (Humphries *et al,* 2015). RIP2 recruitment by activated NOD receptor via CARD-CARD interactions, triggers RIP2 autophosphorylation, filament formation (Pellegrini *et al,* 2018) and both Lys63-Ub and Met1-Ub ubiquitination mediated by different E3 ligases including Inhibitors of Apoptosis (IAPs) and the Linear Ubiquitin Chain Complex (LUBAC) (Boyle *et al,* 2014; Bertrand *et al,* 2009; Damgaard *et al,* 2012; Chin *et al,* 2002; Tao *et al,* 2009; Goncharov *et al,* 2018; Heim *et al,* 2020; Hasegawa *et al,* 2008). Phosphorylation occurs at S176 (Dorsch *et al,* 2006) and Y474 (Tigno-Aranjuez *et al,* 2010), whereas Lys63-Ub has been reported at multiple sites (e.g. K182, K203, K209, K306, K326, K369, K410, K527, K537 and K538) (Hasegawa *et al,* 2008; Goncharov *et al,* 2018; Heim *et al,* 2020). Among the IAPs, the X-chromosome-linked Inhibitor of Apoptosis, XIAP (also known as inhibitor of apoptosis protein 3 (IAP3) and baculoviral IAP repeat-containing protein 4 (BIRC4)) is thought to be the most critical ubiquitin ligase in the NOD2-RIP2 signalling pathway (Krieg *et al,* 2009; Damgaard *et al,* 2012; Bertrand *et al,* 2011; Stafford *et al,* 2018). Similarly to the other IAP proteins, it comprises 3 baculovirus IAP repeats (BIR) domains (BIR1, BIR2 and BIR3) and a C-terminal RING domain (MacE *et al,* 2010). XIAP binds to the kinase domain of RIP2 through its BIR2 domain (Krieg *et al,* 2009; Damgaard *et al,* 2013; Goncharov *et al,* 2018; Hrdinka *et al,* 2018). It then coordinates the conjugation of Lys63-Ub chains (originally thought to be at K209, but now considered to be K410 and K538, see discussion) and triggers the recruitment of LUBAC, which in turn promotes Met1-Ub conjugation and the recruitment of the IKK complex, an essential step for NF-κB activation (Damgaard *et al,* 2012; Hrdinka & Gyrd-Hansen, 2017; Goncharov *et al,* 2018). The absence of XIAP causes defective RIP2 ubiquitination, defective recognition by effector proteins of ubiquitinated RIP2 (e.g. LUBAC), reduced inflammatory signalling and inadequate bacterial clearance (Krieg *et al,* 2009; Bauler *et al,* 2008; Damgaard *et al,* 2012, 2013; Speckmann *et al,* 2013). Mutations in the XIAP gene promote inflammatory pathologies, such as the X-linked lymphoproliferative disease type 2 (XLP-2) and may be related to very early-onset inflammatory bowel disease (Pedersen *et al,* 2014; Damgaard *et al,* 2013; Nielsen & LaCasse, 2017). XLP-2 mutations on the XIAP BIR2 domain abrogate the interaction with RIP2K, and therefore the ubiquitination of RIP2 (Damgaard *et al,* 2013). In the autophagy pathway, NOD2 recruits ATG16L1, which in turn binds the kinase domain of RIP2 and negatively impact its ubiquitination to suppress NF-κB activation by Toll like receptor 2 (Honjo *et al,* 2021a).

RIP2 is an essential downstream component of the NOD and ATG16L1 signalling pathways. RIP2 can also be activated by Toll-like receptor pathways (TLRs), in a way that still needs to be elucidated (Honjo *et al,* 2021b). In all these pathways, recruitment of RIP2 triggers NF-κB activation. Therefore, the inhibition of RIP2 kinase activity has been suggested and demonstrated *in vivo* to be beneficial as a therapeutic strategy for the inflammatory diseases cited above, in particular for IBDs (Honjo *et al,* 2021b; Canning *et al,* 2015; Tigno-Aranjuez *et al,* 2014; Salla *et al,* 2018; Negroni *et al,* 2009; Haile *et al,* 2019; Watanabe *et al,* 2019; Nachbur *et al,* 2015; Philpott *et al,* 2014). However, the molecular significance of the connection between the kinase activity of RIP2, and its role as a scaffold protein for ubiquitination and binding of downstream signalling molecules has remained unclear. Furthermore, several studies (Goncharov *et al,* 2018; Hrdinka *et al,* 2018; Windheim *et al,* 2007; Abbott *et al,* 2004) clearly show that RIP2 auto-phosphorylation activity is not necessary to trigger NF-κB activation, making even more cryptic the role of RIP2 kinase.

In 2018, it was shown that the high nanomolar potency of certain RIP2 kinase inhibitors (e.g. ponatinib, GSK583 and CSLP37/43) depends on these molecules preventing the interaction of RIP2 with the XIAP BIR2 domain and hence inhibiting RIP2 ubiquitination, rather than their ability to inhibit RIP2 kinase activity (Goncharov *et al,* 2018; Hrdinka *et al,* 2018).

In agreement with previous work (Krieg *et al,* 2009; Damgaard *et al,* 2013), Goncharov *et al.* (Goncharov *et al,* 2018) also found that selective XIAP BIR2 antagonists interfere with the XIAP-RIP2 interaction, blocking NOD2-mediated RIP2 ubiquitination and subsequent activation of inflammatory signalling.

Here we present the structure of RIP2 kinase bound to the BIR2 domain of XIAP determined by single-particle cryo-electron microscopy (SPA cryo-EM) at a nominal resolution of 3.15 Å. Combined with biophysical characterisation and *in cellulo* validation of interaction interfaces, the structure provides a molecular explanation for the inhibitory mechanism of antagonists. Moreover, it explains why RIP2K auto-phosphorylation is not required for the protein-protein interaction. The structure is consistent with biochemical data reported in the literature and provides new insight into the regulation of RIP2K ubiquitination.

## Results

### *In vitro* reconstitution of the RIP2K-XIAPBIR2 complex and its stoichiometry

The interaction between RIP2 and XIAP occurs between the kinase domain of RIP2 (RIP2K) and the BIR2 domain of XIAP (Krieg *et al,* 2009; Goncharov *et al,* 2018; Hrdinka *et al,* 2018). Previous interaction studies have used either the XIAP BIR2_AG_^124-260^ (Krieg *et al,* 2009) or the XIAP BIR2_AG_^124-240^ construct (Hrdinka *et al,* 2018; Goncharov *et al,* 2018). Here, the suffix AG denotes two mutations (C202A and C213G) found to improve sample quality for structural studies (Lukacs *et al,* 2013; Sun *et al,* 1999). XIAP BIR2_AG_^124-240^ was used to determine the NMR structure of XIAP BIR2 (Sun *et al,* 1999), whilst the crystal structure was obtained using a shorter construct, XIAP BIR2_AG_^154-240^ (Lukacs *et al,* 2013). Both structures show that XIAP BIR2_AG_^154-240^ is long enough to encompass the folded BIR2 domain (residues 163-230, based on Uniprot annotations P98170), whilst XIAP BIR2_AG_^124-240^ comprises a linker region (residues 124-154), known to be critical to inhibit caspase-3 and −7 activity (Riedl *et al,* 2001; Scott *et al,* 2005; Chai *et al,* 2001).

*In vitro* we reconstituted the RIP2K-XIAP BIR2 complex using both XIAP BIR2^124-240^ and XIAP BIR2^154-240^ as demonstrated by size exclusion chromatography (SEC) and we obtained similar results using the corresponding AG constructs (XIAP BIR2_AG_^124-240^ and XIAP BIR2_AG_^154-240^) (Fig. 1A and 1B, *SI Appendix,* Fig. 1S).

**Figure 1.**
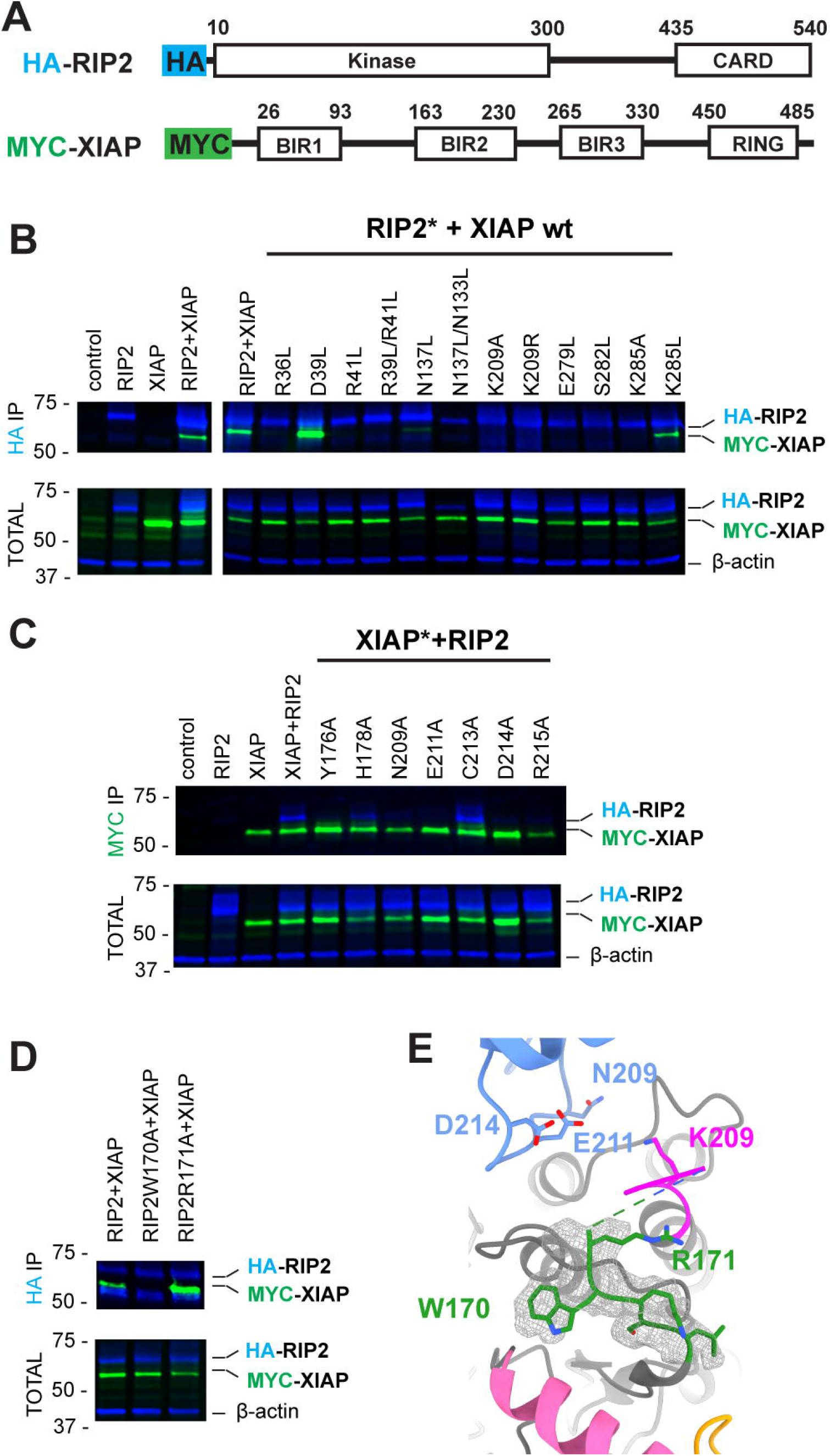
*In vitro* reconstitution and stoichiometry of the RIP2K-XIAP BIR2 complex. (**A**) Overview of the XIAP BIR2 constructs used in this paper. (**B**) SEC profile and SDS PAGE gel of RIP2K^317^-XIAP BIR2^154-240^ complex. (**C**) Native mass spectrometry results for short BIR2 (top spectra) and long BIR2 (bottom spectra) constructs. The main stoichiometry ratios are 2:1 and 2:2 (RIP2K : XIAP BIR2), where two molecules of RIP2K are required for the interaction with XIAP BIR2. The stoichiometry becomes 2:2 in the case of RIP2K-XIAP BIR2_AG_^124-240^.

To determine the stoichiometry of the complexes, we used Native mass spectrometry (MS) on the four samples. When the short BIR2 construct was analysed, the main stoichiometry ratios were 2:1 and 2:2 (RIP2K:XIAP BIR2^154-240^) with higher abundance for the 2:1 oligomer (Fig. 1C, top left panel, *SI Appendix,* Table S1). For the AG construct, the most abundant signals also corresponded to 2:1 and 2:2 ratios of the components (RIP2K:XIAP BIR2_AG_^154-240^), with higher abundance for the 2:2 oligomer (Fig. 1C, top right panel, *SI Appendix,* Table S1). When the longer construct was investigated, the main stoichiometry ratios were also 2:1 and 2:2 for RIP2K-XIAP BIR2^124-240^ (Fig. 1C, bottom left panel). The stoichiometry becomes 2:2 for the RIP2K-XIAP BIR2_AG_^124--240^ sample (Fig. 1C, bottom right panel). These data clearly indicate that two molecules of RIP2K are required for the interaction with XIAP BIR2. Moreover, the data highlight that the AG mutation affects the stoichiometry (*SI Appendix,* Table S1).

### XIAP interacts with RIP2K using exclusively the folded domain

We next investigated whether the linker regions flanking the BIR2 domain contribute to RIP2K binding. We designed several constructs comprising the BIR2 domain (XIAP BIR2^124-263^, XIAP BIR2^93-240^ and XIAP BIR2^93-263^) (Fig.1A) and assessed their ability to bind to RIP2K by determining their dissociation constant (Kd) through Microscale Thermophoresis (MST) measurements. We found that the Kd was in the nM range (Kd= 85.6±18.7 nM) for XIAP BIR2^154-240^ and did not find any significant difference amongst the constructs (*SI Appendix,* Fig. S2).

We then applied solution NMR spectroscopy to define the XIAP BIR2 residues that are involved in the interaction with RIP2K. We successfully produced uniformly ^13^C/^15^N-labelled XIAP BIR2 _AG_ ^124-240^. Backbone (^1^H, ^15^N and ^13^C) NMR assignments of XIAP BIR2 _AG_ ^124-240^ were obtained from a series of three-dimensional HNC-type correlation experiments. Figure 2A shows a superposition of the amide H-N correlation spectra recorded for isolated XIAP BIR2 _AG_^124-240^ (black), and in complex with unlabelled RIP2K (red). Only residues in the N- and C-terminal extension of the XIAP BIR2 construct that are highly flexible in solution remain observable in the complex, while NMR peak intensities for residues in the folded part are significantly attenuated due to the relatively large particle size of the RIP2K-XIAP BIR2 complex (Fig.2B and 2C). These data clearly show that only the folded BIR2 domain is involved in the interaction, while both N- and C-terminal extensions remain highly mobile (Fig. 2B).

**Figure 2.**
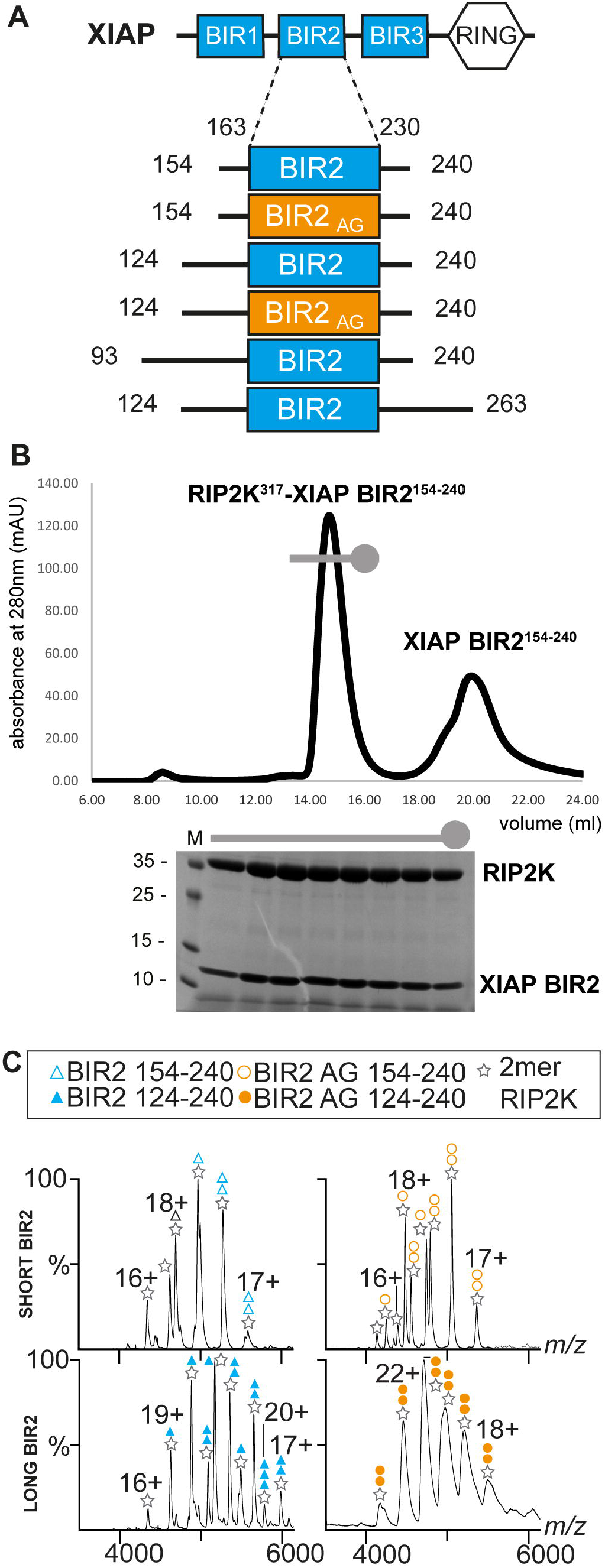
NMR characterization of the XIAP BIR2 interaction with RIP2 in solution. (**A**) Overlap of ^1^H-^15^N BEST-TROSY correlation spectra (25°C) recorded at 850 MHz ^1^H frequency on samples of ^15^N-labeled XIAP BIR2 (black) and a 1 :1 complex of ^15^N-labeled XIAP BIR2 and unlabeled RIP2K (red). BIR2 residues that remain visible in the complex are annotated by their amino-acid type (one-letter code) and residue number. No or only small chemical shift changes are detected for these residues between the free BIR2 protein and the complex, indicating that these protein segments are not involved in the interaction. (**B**) Surface representation of XIAP BIR2 structure (PDB ID: 1C9Q)(56). The N- and C-terminal protein segments that are not involved in the interaction with RIP2 are highlighted in red. (**C**) {H}-N heteronuclear NOE (HETNOE) ratios measured for XIAP-BIR2 at 25°C and 850 MHz ^1^H frequency. The protein segments that remain visible in the NMR correlation spectrum of (A) upon interaction with RIP2K are highlighted by red bars. These N- and C-terminal segments show reduced HETNOE ratios (≤ 0.6), indicative of significant fast (sub-ns) time scale backbone mobility, while for the central part an average HETNOE of 0.8 is measured, in agreement with a globular protein domain. The flexibility of the N- and C-terminal segments is preserved in the complex, which makes them NMR-observable despite its high molecular weight.

Furthermore, NMR-detected translational diffusion measurements indicate that upon interaction with RIP2 the apparent molecular size is increased by a factor of about 5.7 *(SI Appendix,* Fig. S3). This is in good agreement with the presence of a RIP2 dimer in the complex, and a 1:2 BIR2:RIP2 stoichiometry, although the presence of 2:2 BIR2:RIP2 complexes cannot be excluded. Based on the results obtained so far, we decided to use only the short BIR2 constructs (residues 154-240) for structural studies.

### Structure determination by cryo-EM of RIP2K-XIAP BIR2 complex

Exhaustive crystallisation trials failed to produce crystals containing either the RIP2K-XIAP BIR2 or the RIP2K-XIAP BIR2_AG_ complex, resulting only in RIP2K dimer crystals. Therefore, we attempted to solve the structure of RIP2K-XIAP BIR2 by cryo-EM. To make the complex, we purified both wild-type domains to homogeneity and combined them in the presence of ATP-MgCl_2_ with an excess of XIAP BIR2 protein. The sample was further purified by size-exclusion chromatography and the most enriched complex fraction was selected for vitrification (*SI Appendix,* Fig. S4). We performed data collections with a Titan Krios, combining untilted and tilted data collections. We obtained two-dimensional (2D) class averages showing structural features and finally obtained a map at a nominal resolution of 3.15 Å (*SI Appendix,* Fig. S4, Fig. S5, Fig.S6 and Table S2). In this map, we were able to unambiguously dock two RIP2K molecules, arranged as an antiparallel dimer, as in RIP2K crystal structures (PDB ID: 5NG0, 4C8B), bound with one molecule of XIAP BIR2 (PDB ID:4J3Y) (Pellegrini *et al,* 2017; Lukacs *et al,* 2013; Canning *et al,* 2015) (Fig.3 and *SI Appendix,* Fig. S7, Table S2). We immediately observed that one of the BIR2 mutations (the C213G) known to improve protein quality, is at the interaction interface (Fig. 3F). Therefore, we did not pursue the structure determination of the RIP2K-XIAP BIR2_AG_ complex.

**Figure 3.**
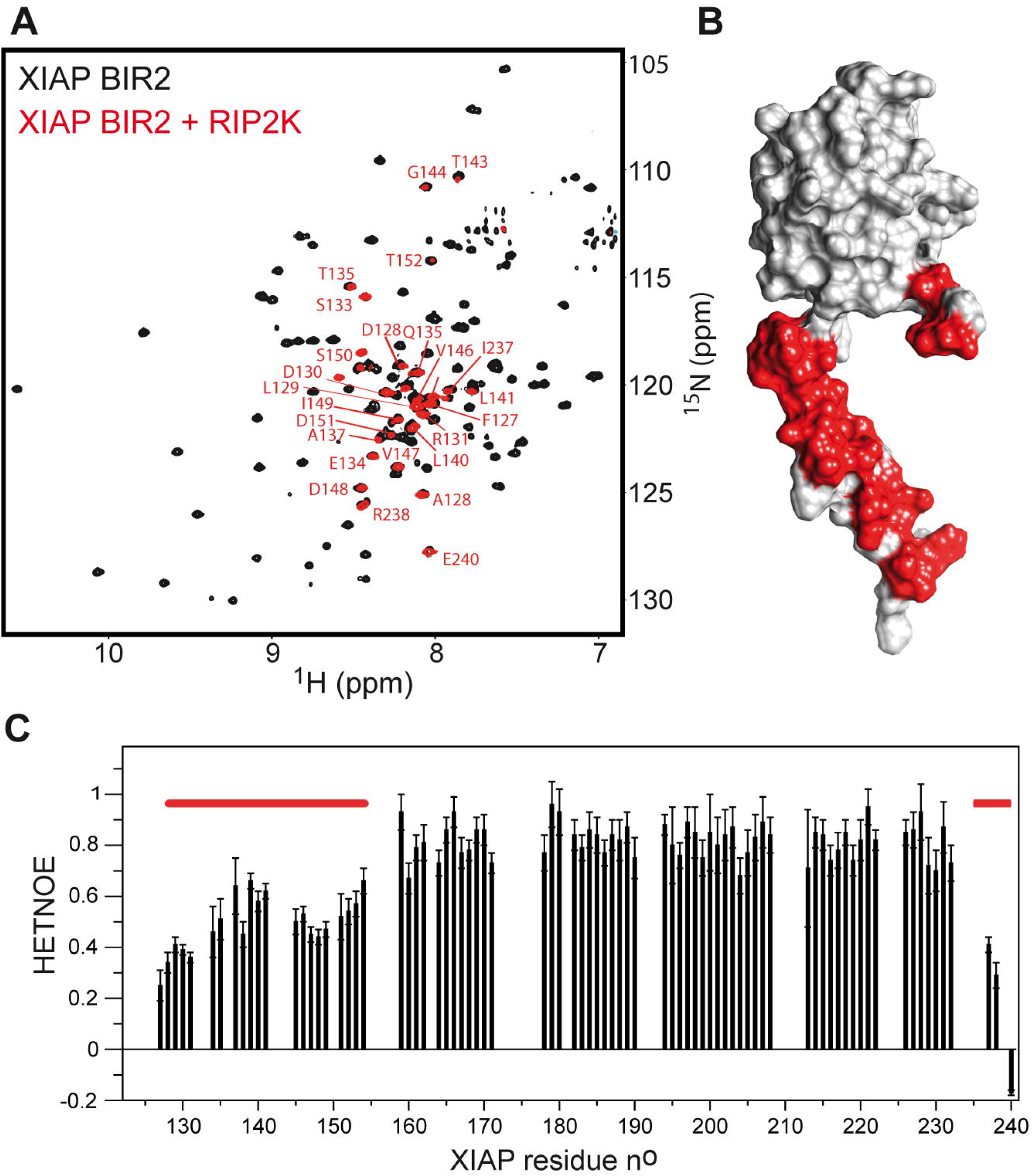
Structure of the RIP2K-XIAP BIR2 complex. (**A**) Ribbon representation of the RIP2K^317^-XIAP BIR2^154-240^ structure. XIAP BIR2 is colored in light blue, kinases molecules in light and dark grey (Kinase_A, Kinase_B), αC-helix in pink (residues 52-72), Gly-rich loop in yellow, activation loop in green and K209 loop in magenta. (**B-F**) Assignment according to the cryo-EM density features of (**B**) Kinase_A αE-helix residues (120-137), (**C**) Kinase_B αE-helix residues (120-137), (**D**) Kinase_A and Kinase_B αL-helices and their interaction (residues 299-312), (**E**) XIAP BIR2 interaction with Kinase_A, (**F**) XIAP BIR2 interaction with Kinase_B.

Both kinase chains are in the active-like conformation based on their αC-helix position (IN conformation) and density can be observed for nucleotide in its active site (Fig. 3A and *SI Appendix* Fig.S7). The N-termini anti-parallel β-strands described to be present in the dimeric structure of active-like RIP2K (Pellegrini *et al,* 2017), is absent (*SI Appendix* Fig.S8).

In parallel, we used Alphafold2 as implemented in Colabfold (Mirdita *et al,* 2022; Jumper *et al,* 2021; Evans *et al,* 2022) to predict the structure of dimeric RIP2K with bound XIAP BIR2. In agreement with our experimentally derived model, the predication shows no anti-parallel β-strand interaction of the kinase N-termini. Interestingly, the Predicted Aligned Error diagram (PAE) show low confidence interaction between the kinase and the BIR2 domain, despite the predicted model being very close to that observed (*SI Appendix* Fig.S8C).

### The RIP2-XIAP interaction requires RIP2K dimerization

The final model of the complex corresponds to a mass of 84 kDa (*SI Appendix* Table S1), with a 2:1 stoichiometry, in agreement with both Native mass spectrometry and solution NMR data. Only one molecule of XIAP BIR2 is bound at the dimer interface and it interacts with the C-lobe of one kinase molecule (Kinase_A) and the N-lobe of the other one (Kinase_B) (Fig. 3A). XIAP BIR2 uses loop 176-178 to mainly interact with Kinase_A and loop 209-214 to interact with Kinase_B (Fig. 3E and 3F). To confirm observed interactions, we mutated to either alanine or leucine the residues at the XIAP BIR2-RIP2K interaction interface. The mutations were performed in tagged full length constructs, which were transiently expressed in mammalian cells, and pull down assays were used to assess the effect of the mutations on complex formation (Fig. 4).

**Figure 4.**
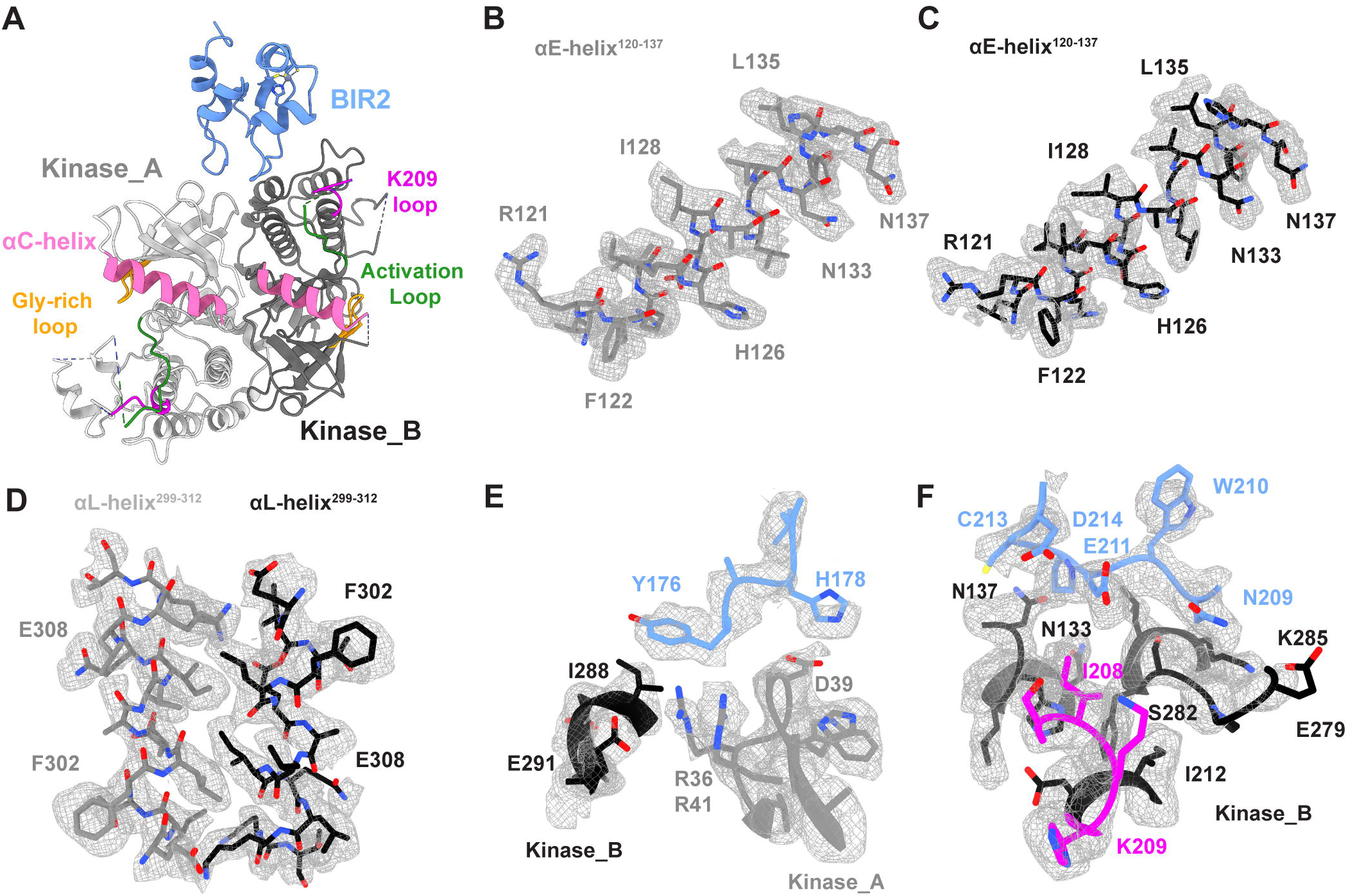
Validation of the interaction interfaces between RIP2K and XIAP BIR2. (**A**) Schematic representation of the RIP2 and XIAP constructs used for expression of wt and mutants proteins in HEKT293 cells. (**B-D**) Results of expression and pull down (IP, immunoprecipitation) on HA-RIP2 and MYC-XIAP proteins from HEKT293 cells. (**E**) Molecular environment of the Kinase_B activation loop (L164-R171) and its architecture in the density

XIAP BIR2 loop 176-178 comprising Y176 and H178, makes electrostatic interaction with a hydrophilic patch formed by the loop between strands β2 and β3. Mutations of XIAP Y176A and mutations in the RIP2-Kinase_A interaction interface (R36L, R41L, R36L/R41L) destroy the interactions. XIAP H178 potentially make hydrogen bonds with the oxygen group of D38. However, the mutations XIAP H178A and RIP2 D39L do not impair complex formation (Fig. 3E, 4B and 4C).

The XIAP BIR2 loop 209-214 makes electrostatic interactions with the C-lobe of Kinase_B. N209, E211 and D214 form a negatively charged binding site that appears to lock Kinase_B residues K209-I208. Mutations of these XIAP loop residues block complex binding, except for C213A, which agrees with the fact that the XIAP BIR2_AG_ construct can still bind the kinase. Mutations of RIP2 K209, E279, S282 and K285 destroy the binding, validating the interaction interface. Moreover, double mutations N137L/ N133L also prevents binding (Fig. 3F, 4B and 4C).

The activation loop of Kinase_B is mostly disordered and XIAP BIR2 does not interact with the visible part (residues 164-171). However, we found that mutation at W170 but not R171 impair complex formation (Fig.4D and 4E).

## Discussion

In crystal structures, RIP2K is always an antiparallel dimer (Canning *et al,* 2015; Charnley *et al,* 2015; Pellegrini *et al,* 2017). We previously showed that RIP2K is also in a dimeric state in solution suggesting that kinase activation is coupled to dimerization (Pellegrini *et al,* 2017). This is also true for the D146N mutant, which corresponds to a protein without kinase activity, yet still able to trigger NF-κB activation (Goncharov *et al,* 2018; Hrdinka *et al,* 2018). In the case of the K47R (or K47A) dead kinase, NF-κB activation is still possible and the protein is both dimeric and monomeric in solution (Pellegrini *et al,* 2017; Hrdinka *et al,* 2018; Goncharov *et al,* 2018). Hitherto, a rationale for the RIP2 dimer in NOD signalling has not been identified.

The reconstruction of the RIP2K-XIAP BIR2 complex shows the kinase in a similar antiparallel dimer as in the RIP2K crystal structures. Consistent with our NMR results, only the folded BIR2 domain binds to RIP2K. Importantly, the bound XIAP BIR2 domain engages both monomers, highlighting the importance of the RIP2K dimer as scaffold for downstream ubiquitination. Binding is driven by the XIAP BIR2 loops and the resulting interface is different from the one described for the caspase 3 and 7 binding sites (PDB Ids: 1I3O,1l51 and *SI Appendix* Fig.S9A and 9B) (Riedl *et al,* 2001; Chai *et al,* 2001). All the residues previously described to affect to the RIP2K-XIAP BIR2 interaction contribute directly to protein-protein interaction, except I212, which is located deeper in the Kinase B structure (Damgaard *et al,* 2013; Hrdinka & Gyrd-Hansen, 2017; Cavallari *et al,* 2017). In particular, the two RIP2K patches suggested to be part of the interaction, the R36-R41 basic patch (Hrdinka *et al,* 2018) and the I208-K209 patch (Heim *et al,* 2020), which are 40Å apart in the monomer, are in fact close to each other in the dimer because of the antiparallel kinase arrangement.

No interaction is observed between the XIAP BIR2 domain and the kinase activation loop, which indicates that XIAP binding requires RIP2K to be in dimeric and αC-helix IN conformation, but not necessarily phosphorylated at the activation loop. This agrees with the fact that wild-type and D146N mutant both form the same antiparallel dimer, the K47R mutant can still form it in solution and all three constructs trigger NF-κB signalling (Goncharov *et al,* 2018; Hrdinka *et al,* 2018). Our structure supports that the phosphorylation state is not relevant for NF-κB activation in the NOD signalling.

Native mass spectrometry data shows both 2:1 and 2:2 stoichiometry, whereas NMR translational diffusion measurements suggest a 2:1 stoichiometry. We found these data quite surprising, as we expected two XIAP BIR2 binding sites based on the symmetrical arrangement of the kinase dimer. In agreement with the biophysical data, we could not identify any 2D classes with double XIAP BIR2, suggesting that the presence of a second copy of XIAP BIR2 in the RIP2K-XIAP BIR2_AG_^124-240^ sample might be related to the ability of XIAP BIR2 domain to homodimerise (Lukacs *et al,* 2013). Interestingly, the complex reconstruction does not show any antiparallel β-sheet interaction of the kinase N-termini, which appear to be flexible. Moreover, the two kinase monomers are closer in comparison to the original active structure (PDB ID: 5NG0, *SI Appendix* Fig.S8) (Pellegrini *et al,* 2017). From these observations, we speculate that binding of XIAP BIR2 perturbs the kinase dimer, which might be the reason for which only one XIAP BIR2 molecule can be bound.

Based on the RIP2K-XIAP BIR2 structure, we can now provide a molecular explanation for the mechanism of XIAP BIR2 and RIP2K inhibitors that prevent complex formation. Several papers reported that SMAC mimetic compounds (SMCs), which bind in the so called BIR IBM (IAP-binding motif) binding groove, abolish the XIAP BIR2 interaction with RIP2 resulting in impaired ubiquitination of RIP2 protein and diminished NF-κB signalling (Krieg *et al,* 2009; Damgaard *et al,* 2013; Goncharov *et al,* 2018). These compounds mimic the IBM, a 4 residue linear motif (AVPI) belonging to the SMAC/DIABLO protein (second mitochondria-derived activator of caspases/direct IAP-binding protein with low pI), which antagonises the binding of caspases to IAP proteins (Gyrd-Hansen & Meier, 2010). The XIAP BIR2 domain binds to the IBM of activated caspase-3 and caspase-7 (Scott *et al,* 2005). Comparison between our structure and that of XIAP BIR2 in complex with peptide AVPI (PDB ID: 4J46,)(Lukacs *et al,* 2013), clearly shows that BIR2 inhibitor prevents complex formation with RIP2K occluding the IBM groove (*SI Appendix* Fig.S9C). This is in agreement with the fact that X-linked lymphoproliferative disease type 2 (XLP-2) BIR2 mutations that either destabilise the domain or belong to the IBM groove, abolish the RIP2K-BIR2 interaction (Lukacs *et al,* 2013; Sun *et al,* 1999; Damgaard *et al,* 2013).

Hrdinka *et al.* observed that RIP2K inhibitors that affect both kinase activity and ubiquitination state (e.g. Ponatinib, GSK583, CSLP37/ 43 and 7f) interact with residues of the so called back pocket of the kinase (A45, K47, I96 and gatekeeper T95), a binding site that becomes accessible when the activation loop is in the inactive conformation. They further speculated that binding of inhibitors to this pocket could induce conformational changes at the basic patch-surface exposed on the top of it, comprising R36 and R41. Single and double mutations of these two arginines impairs the interaction with XIAP-BIR2 domain (Fig.4B), (Hrdinka *et al,* 2018). Our structure shows that these residues are involved in the RIP2K-BIR2 interaction, and they are positioned at the kinase dimerization interface (Fig. 3E). Indeed, any perturbation of this zone might prevent binding between the two proteins by affecting either the binding itself or by destabilising the kinase dimer.

Crystal structures of RIP2K in complex with Ponatinib, GSK583 or CSLP18, show that inhibitor binding perturbs the DFG-in conformation and disorders the activation loop beyond residue 164, whilst in the active conformation residues 164-171 can still be traced (PDB IDs: 4C8B, 5J7B and 6FU5, *SI Appendix* Fig.S10) (Canning *et al,* 2015; Haile *et al,* 2016; He *et al,* 2017). Therefore, we used our *in cellulo* pull down assay to test whether the conformation of the visible part of the activation loop is important for the interaction. We mutated W170 and R171 to alanine, and found that the W170A mutation abolishes complex formation, suggesting that the conformation of the activation loop up to W170 is important (Fig. 4D and 4E). Therefore, the ability of inhibitors to completely destabilise the activation loop and the DFG-in conformation might contribute to their capacity to prevent complex formation.

Interestingly, the RIP2K-XIAP BIR2 structure shows that RIP2 K209 residue is part of the protein-protein interface (Fig.3F). K209 was originally proposed as a ubiquitination site, based on the observation that the K209R mutant blocks RIP2K ubiquitination and it has a loss of function phenotype when overexpressed with ubiquitin (Hasegawa *et al,* 2008; Tigno-Aranjuez *et al,* 2013). However two recent studies, which used a proteomic approach to determine RIP2 ubiquitination sites by XIAP, failed to detect ubiquitination at K209 and instead suggested that the relevant lysines were K410 and K538 (Goncharov *et al,* 2018; Heim *et al,* 2020). Moreover, our pull down data, in agreement with published data (Heim *et al,* 2020) show that mutants K209A and K209R block the RIP2K-BIR2 interaction (Fig.4B).

Alternatively, K209 could be a ubiquitination site targeted by other IAPs, such as cIAP1 and 2 (Bertrand *et al,* 2009, 2011), even though it is still controversial whether these cIAPs contribute to NOD signalling by ubiquitination of RIP2 (Stafford *et al,* 2018). BIR domains shared a common fold, and their ability to interact with different partners depends on sequence differences at surface exposed positions (Budhidarmo & Day, 2015). Comparison of the crystallographic structure of XIAP BIR2 domain with the cIAP1 and 2 BIR2 domain predicted using Alphafold2 shows high structural conservation, for the IBM groove (*SI Appendix* Fig.S11). Indeed sequences of the BIR2 domains of XIAP, cIAP1 and cIAP2 BIR2 show high conservation in the XIAP loop 200-214 (*SI Appendix* Fig.S11A). Our pulldown assay shows that single mutants of XIAP residues N209, E211 and D214 in loop 200-214 abolish protein-protein interaction (Fig.4B). These observations suggest that the BIR2 domains of XIAP and cIAP1-2 are highly similar and they probably interact with RIP2K dimer in a similar fashion, making inaccessible K209 for ubiquitination. Our results provide strong evidence that the importance of K209 in RIP2 ubiquitination, is related to its fundamental contribution to the binding of BIR2.

## Conclusion

The biophysical and structural characterisation of the RIP2K-XIAP BIR2 interaction described here reveals that the scaffold function of RIP2K in binding XIAP requires it to be a dimer. This study also provides a molecular explanation why the phosphorylation state of RIP2K is not relevant for NOD signalling and why XIAP BIR2 antagonists and potent RIP2K inhibitors abrogate RIP2 ubiquitination and NF-κB activation by preventing complex formation. Moreover, considering the RIP2K-XIAP BIR2 structure together with our *in cellulo* validation and previously published observations, we confirm that K209 is critical for RIP2 ubiquitination most likely because it contributes to the RIP2K-XIAP BIR2 interaction interface. In conclusion, this study, by providing explanations for several previously suggested mechanisms, advances our knowledge on the scaffold role of the RIP2 protein and its interaction with XIAP BIR2. These results might be of value for the design of more potent and specific RIP2K and XIAP BIR2 inhibitors.

## Methods

### Protein constructs

Recombinant human RIP2K ^1-300^ and RIP2K ^1-317^ were produced using the baculovirus system in *sf21* insect cells. Cloning of DNA encoding RIP2K^1-300^ with a Tobacco Etch virus protease (TEV) cleavable maltose binding protein tag (MBP-tag) at the N-terminus in pFastBacHTB has been previously described (Pellegrini *et al,* 2017). Extension to residue 317 (RIP2K^1-317^) of wild-type construct has been added by using the ABC (restriction/ligase) free method (Qaidi & Hardwidge, 2019).

Plasmids pET49 encoding for human XIAP BIR2^154-240^ and XIAP BIR2_AG_^154-240^ are a gift from Katrin Rittinger (Francis Crick Institute, UK). Both proteins comprise a Prescission protease (P3C) cleavable Glutathione-S-transferase (GST-tag) at the N-terminus. Constructs XIAP BIR2^124-240^, XIAP BIR2_AG_^154-240^, XIAP BIR2^124-263^ XIAP BIR2^93-240^ and XIAP BIR2^93-263^ were cloned using the ABC method. The cloning resulted in 2 additional residues (GP) following the P3C cleavage site.

The sequences of the oligomers used for constructs extension are reported in *SI Appendix* Table 3.

### Protein expression and purification

N-terminally HIS-tagged TEV and N-terminally GST-tagged P3C proteases used for protein purification were produced at the Protein Expression and Purification Core Facility at EMBL, Heidelberg, Germany.

RIP2K and RIP2K were expressed and purified using a similar protocol (Pellegrini *et al,* 2017). Bacmid generation, transfection, virus production, virus amplification and protein expression have been performed at the EMBL Eukaryotic Expression Facility, following guidelines provided by the facility. Proteins have been expressed in *sf21* cells, using Sf-900 SFM medium (Gibco Life technologies). Cells were harvested 4 days post infection and were lysed by sonication in buffer A (20 mM Tris pH 7.5, 300 mM NaCl, 50 mM NDSB, 5% glycerol, 500 μM TCEP) containing protease cocktail inhibitor (Complete, Roche). After centrifugation at 20,000 ×g for 30 min at 4□°C, the supernatant solution was incubated for at least 2□h with amylose-affinity chromatography resin (New England Biolabs), whilst gently shaking at 4□°C. The fusion protein was then eluted using the same lysis buffer supplemented with 40 mM maltose. Upon overnight TEV cleavage and dialysis against buffer B (20 mM Tris pH 7.5, 50 mM NaCl, 50 mM NDSB, 5% glycerol and 500 μM TCEP), proteins were further purified by anion exchange chromatography with a 0 to 1M NaCl gradient, over two column volumes. Fractions corresponding to pure RIP2K were then pooled and applied onto a PD-10 desalting column (GE Healthcare) equilibrated with buffer C (20 mM Tris pH 7.5, 150 mM NaCl, 2% glycerol). Proteins were then concentrated to 1-1.5 mg/ml and frozen in liquid N_2_ for storage at −80 °C.

For *in vitro* reconstitution of the complex and Native MS experiments, both RIP2K_1-300_ and RIP2K_1-317_ were used. For NMR and cryo-EM application, we exclusively used RIP2K_1-317_.

XIAP BIR2 constructs were expressed in *E. coli* Rosetta 2 (Novagen) by growing the bacterial culture at 37 °C until an OD_600□nm_ of 0.6 and inducing with 0.250 mM IPTG (isopropyl-β-D-1-thiogalactopyranoside) overnight at 16 °C. Cells were harvested and re-suspended in buffer A supplemented with 10 μM ZnCl_2_ and protease inhibitor (Complete, Roche). Cells were then homogenized by sonication and the crude extract was centrifuged for 30 min at 18,000□×□*g.* Clear lysate was incubated with Glutathione sepharose resin (BRAND) for 2 hours, whilst gently shaking at 4□°C. The beads were then washed with buffer A and the protein was eluted in buffer A supplemented with 20 mM reduced L-Glutathione. Upon overnight P3C cleavage and dialysis against buffer D (20 mM Tris pH 7.5, 150 mM NaCl, 25 mM NDSB, 5% glycerol, 500 μM TCEP), the cleaved protein was separated from un-cleaved protein, GST and GST-P3C by repeating the affinity step without incubation. Eluate was then aliquoted and frozen in liquid N_2_ for storage at −80 °C.

### Microscale Thermophoresis (MST)

As Tris is not compatible with primary amine fluorescent labeling, buffer for all the samples (RIP2K^1-300^, XIAP BIR2^154-240^, XIAP BIR2^124-240^, XIAP BIR2^93-240^ and XIAP BIR2^93-263^) was exchange by SEC in buffer E (20 mM Hepes pH 7.5, 150 mM NaCl, 2% glycerol). Labeling of RIP2K^1-300^ was performed with Monolith Protein labeling kit RED-NHS 2nd generation (MO-L011, Nanotemper) following the manufacturer’s instructions. For labeling, the protein concentration was adjusted to be in the 6-10 μM range. For measurements, labeled RIP2K^1-300^ was diluted to 40 nM, whilst the XIAP BIR2 constructs were titrated from 1,2 to 2500 nM. 10 μL of each XIAP BIR2 constructs were mixed with 10 μL of labeled RIP2K^1-300^ at final concentration of 20 nM. Glass capillaries were filled with 4μL of each mixture. Signal was improved by adding 0,05 % Tween-20, by choosing red fluorescent excitation wavelength and by setting the LED power at 50%. All measurements were performed at 20 °C.

### NMR

For NMR measurements, two samples were prepared: N, C-labelled XIAP BIR2_AG_ ^124-240^ and labelled XIAP BIR2_AG_^124-240^ in complex with RIP2K^1-317^.

XIAP BIR2_AG_^124-240^ was expressed in *E. coli* Rosetta 2 (Novagen) by growing the bacterial culture at 37 °C until an OD_600□nm_ of 0.6 and inducing with 0.500 mM IPTG for 5 hours at 30°C in minimal growth medium M9 supplemented with either ^15^N-labeled NH_4_Cl or a combination of^15^NH4Cl and ^13^C-labeled glucose. Labelled XIAP BIR2_AG_^124-240^ (^15^N-XIAP BIR2_AG_^124-240^ and (^15^N ^13^C) XIAP BIR2_AG_^124-240^) were purified as described above. After the second affinity step, samples were applied onto a Superdex 200 (10/300) increase (GE Healthcare) column equilibrated in buffer F (20 mM Tris pH 7.5, 150 mM NaCl). Labelled XIAP BIR2_AG_^124-240^ was directly concentrated on 3 KDa molecular weight cut off Centricon centrifugal filter units (EMD Millipore) until 300 μL at 240 μM were obtained.

RIP2K^1-317^ buffer exchange in buffer F was done by using a PD-10 desalting column (GE healthcare). 5mM MgCL_2_ and 2 mM ATP were added to the kinase sample, and labelled XIAP BIR2_AG_^124-240^ was added in ratio 1.1:1 (RIP2K:XIAP BIR2). Sample was let on ice for 30 min then concentrated on 3 kDa molecular weight cut off Centricon centrifugal filter units (EMD Millipore) until 300 μL at 200 μM were obtained.

All NMR experiments were performed at 25°C on Bruker Avance IIIHD spectrometers operating at magnetic field strengths of 850 or 950 MHz (H frequency), equipped with cryogenically cooled triple-resonance probes. H-N correlation spectra were recorded using a BEST-TROSY pulse scheme (Favier *et al,* 2011), while chemical shift assignments were obtained from a set of 3D BEST-TROSY HNC correlation experiments (Solyom *et al,* 2013). Translational diffusion constants of the proteins in solution were measured by 1D H DOSY experiments (Johnson, 1999). NMR data processing and analysis was performed using TopSpin 3.5 (Bruker BioSpin) and CCPNMR V3 software tools.

### Complex purification for Native MS and cryo-EM

In order to remove residual GST and protein aggregates, thawed XIAP BIR2 sample was applied onto a Superdex 200 (10/300) increase (GE Healthcare) column equilibrated in buffer C. In parallel, thawed RIP2K was incubated with 500 μM AMPPCP and 5 mM MgCl_2_ and let on ice for 30 min. Homogeneous XIAP BIR2 was then added in excess (ratio: 1:1.3) and complex (RIP2K^1-317^ with either XIAP BIR2^154-240^ or XIAP BIR2_AG_^154-240^) was let 30 min on ice. Sample was then concentrated 6 times by ultrafiltration using a 3 kDa molecular weight cut off Centricon centrifugal filter units (EMD Millipore) and applied onto a Superdex 200 10/300 increase (GE Healthcare) in buffer C to remove excess of XIAP BIR2. Homogeneity of the sample was then check by SDS PAGE gel (Novagen), and the most concentrated complex fraction, usually in the concentration range of 5-7 μM, was used for cryo-EM grids preparation without further dilution. For Native MS experiments, fractions containing the complex were re-applied onto a Superdex 200 10/300 increase equilibrated in buffer G (250 mM ammonium acetate with 1 mM DTT) and measurements were immediately run afterwards.

### Native MS

RIP2K^1-300^ or RIP2K1^-317^ in complex with either XIAP BIR2^154-240^ or XIAP BIR2_AG_^154-240^ were analysed by Native MS in the concentration range of 10 μM. Measurements were run as previously described (Pellegrini *et al,* 2017).

### Cryo-EM specimen preparation

Cryo-EM specimens were prepared on UltrAuFoil R1.2/1.3 holey gold grids (Quantifoil) that were glow-discharged for 20 s at 25 mA on both sides (PELCO easy glow). Vitrobot (Vitrobot Mk IV, Thermo Fisher Scientific) was set to 4 °C and 100% humidity. A drop of 2 μl sample was applied to each grid side and blotting was run at force 0, for a total time of 3.5-4.5 s. Grids were then vitrified by plunging into liquid ethane at liquid N2 temperature. Grids were clipped into autoloader cartridges and screened using a Glacios cryo electron microscope (Thermo Fisher Scientific) equipped with a Falcon 3 detector. Promising grids showing visible particles by eye at −1 μM defocus, were used for data collection on CM01, ESRF Grenoble France (*SI Appendix* Fig.S4).

### Cryo-EM Data Collection and Processing

The RIP2K^1-317^ -XIAP BIR2^154-240^ dataset was acquired using a Titan Krios operating at 300 keV, equipped with K2 Quantum detector (Gatan) and a GIF Quantum energy filter (Gatan) at CM01 (ESRF, Grenoble, France)(Kandiah *et al,* 2019). 7178 movies were collected in untilted mode, at 165K magnification, corresponding to a pixel size of 0.87 px/Å for a total dose of 47.8 (1.2 e-/A2 per frame, fractionated in 40 frames). A total of 3698 movies were recorded in tilted mode (1870 micrographs at 20 degrees, 1825 micrographs at 25 degrees), at the same magnification, for a total dose of 45.7 (1.1 e-/A2 per frame, fractionated in 40 frames) (*SI Appendix* Fig.S4, Table S2).

Movies were imported into Relion, aligned and dose-weighted using MotionCor2 in Relion 3.1 (Zheng *et al,* 2017; Fernandez-Leiro & Scheres, 2017). Micrographs were then imported into CryoSPARC (Punjani *et al,* 2017) were CTF estimation has been run with Patch CTF and tilted micrographs were manually curated according to CTF based estimated resolution (<10Å) and estimated ice thickness (<1.1). Any attempts of processing of the untilted dataset alone or in combination with the tilted datasets with a series of 2D and 3D classifications, resulted in poorly resolved cryo-EM maps indicating preferred particle orientations. Therefore, we modified our processing approach. We firstly processed only the tilted data. The two tilted datasets (collected at 20 degrees and 25 degrees respectively) were kept separately and processed similarly (*SI Appendix* Fig.S5). Particles were firstly picked on 100 representative micrographs using Topaz (Bepler *et al,* 2019) and they were extracted with a box size of 300 x 300 pixels. We applied 2D classification with a tight mask (100 Å) and select 2D-classes showing particle features. These particles were used to train Topaz and the resulting topaz_model was used to pick particles in the remaining micrographs. After extraction, 2D classification was applied to eliminate bad particles and to select 2D class averages with lower background noise and stronger features. We then performed *ab-initio* reconstruction with two models followed by non-uniform (NU) refinement using the best model as reference and all the particles. The resulting map, denoted the tilted map, showed a better orientation distribution than for previous maps. In order to maintain a balance between the number of particles from the tilted dataset and the number of particles from the untilted dataset, only 2000 micrographs of the later were selected. Particles were then picked using as template projections of the tilted map. Resulting particles were extracted and submitted to 2D classification to eliminate bad particles. Remaining particles were then combined with tilted ones, and submitted to another cycle of 2D classification. We then selected the particles with high effective classes assigned values (ECA, between 2.1 and 2.6), as in our case these were the ones showing the sharpest features and the lowest background noise. From the NU-3D refinement of these particles, we obtained the final map, at an average resolution of 3.15 Å (FSC 0.143 threshold). This map was used to calculate directional FSC and local resolution in CryoSPARC (*SI Appendix* Fig.S6). Adding more particles did not improve the map resolution and promoted anisotropic resolution. Further 3D classification using either heterogeneous refinement or 3D classification (beta) with different masks size did not improve particle selection nor the clarity of final map. A processing workflow, comprising data statistics, is shown in *SI Appendix* Fig.S5.

For map sharpening, we used the sharpening tool in CryoSPARC. For manual reconstruction in Coot, the map was sharpened and blurred using the mrc_to_mtz tool in ccpem (Wood *et al,* 2015; Burnley *et al,* 2017).

### Model building and refinement

The atomic model of RIP2K^1-317^ in complex with XIAP BIR2^154-240^ was obtained by fitting with rigid-body refinement the existing crystal structures of both proteins. For RIP2K, we firstly fit the structure of active RIP2K (PDB ID: 5NG0)(Pellegrini *et al,* 2017) which displays RIP2K residues from 5 to 310. For XIAP BIR2, we used the crystal structure 4J3Y, chain C (Lukacs *et al,* 2013). We firstly fit the two structures in the cryo-EM map, using ChimeraX (Pettersen *et al,* 2021). After removal of waters, ions and nucleotide analogue, we then proceeded with rigid-body refinement in Phenix (Liebschner *et al,* 2019). In Coot, we mutated back to cysteine residues 202 and 212 in the BIR2 domain, manually adjusted loop 210-214, built the αL-helices (residues 299-312) using as template the structure of RIP2K-ponatininb (PDB ID: 4C8B)(Canning *et al,* 2015), and deleted the regions obviously outside of the density as well as the N-termini β-strand interaction. We then applied a second cycle of rigid-body refinement to obtain the final model. Figures were prepared with ChimeraX 1.4 (Pettersen *et al,* 2021). Software used in this project was installed and configured by SBGrid (Morin *et al,* 2013). Refinement statistics are reported in *SI Appendix* Table S2.

### Alphafold2

For the Alphafold2 calculations we used Local ColabFold (Evans *et al,* 2022; Jumper *et al,* 2021; Mirdita *et al,* 2022) installed on an in house server. To compute the complex structure of XIAP BIR2 with RIP2K (residues 154-240, 1-317 respectively) we used 3 cycles, model ‘multimer-2’ and 5 models. Neither increasing the number of cycles nor changing module improved the prediction. We applied the same calculation to RIP2K-cIAP1 BIR2 and RIP2K-cIAP2 (*SI Appendix* Fig.S11C). As for RIP2K-XIAP BIR2 complex, the prediction does not show any significant interaction between the two proteins.

### Mammalian cell culture and plasmids

A HEK293T cell line (from the laboratory of W. Filipowicz) was used, as previously described (Pellegrini *et al,* 2018). Cells were maintained in DMEM medium (Lonza) supplemented with 10% (v/v) foetal bovine serum (FBS) and nonessential amino acids (Gibco), at 37 °C and 5% CO_2_. Human XIAP full length (1-497) construct for *in cellulo* experiments, was purchased (puno1-hxiap, InvivoGen) and it was cloned from pUNO1 into the vector pcDNA3 using BamHI and XhoI restriction sites. Using the ABC method (Qaidi & Hardwidge, 2019) a P3C cleavable MYC tag was added at the N-terminus together with a linker (ASASAS), resulting in pcDNA3-MYC-XIAP(1-497). Used oligomers are reported in *SI Appendix* Table S3.

Single amino acids mutants (RIP2: R36L, D39L, R41L, R39L/R41L, N137L, N137L/N133L, K209A, K209R, E279L, S282L, K285A and K285L; XIAP: Y176A, H128A, N209A, E211A, C213A, D214A and R215A) were obtained by site directed PCR mutagenesis of pcDNA3-HA-RIP2(1–540), and pcDNA3-MYC-XIAP (1-497) using the oligomers listed in *SI Appendix* Table S3.

### Immunoprecipitation and Western blot

For co-immunoprecipitation of MYC-XIAP, HA-RIP2 and relative mutants, HEK293T cells were seeded in 6-well plates, 24 h prior transfection. Transfection was performed with LipoD293 transfection reagent (SignaGen). Each well was transfected with 275 ng of either pcDNA3-HA-RIP2 or corresponding mutant and 725 ng of either pcDNA3-Myc-XIAP or corresponding mutant. As negative control, a well was transfected with 1000 ng of pcDNA3 empty vector. Each transfection mixture was completed with NOD2 activator MDP (tlrl-mdp, InvivoGen) and cells were lysed 22h after transfection in 143 μl of Cell Lysis buffer (Cell Signaling) supplemented with protease inhibitor (Complete, Roche). According to the manufacturer’s protocol, after 5 min incubation on ice, cells were scratched from each well, and sonicated briefly (2 cycles of 7s intercalated by incubation on ice). Residual cell debris were eliminated by centrifuging at 13 000 g for 15 minutes. 400 μl of clean sample was then incubated overnight at 4 °C on gentle shaking with 20 μl of either anti-MYC-agarose beads (3400, Cell Signaling) or anti-HA-agarose beads (3956, Cell Signaling). Beads were then washed 5 times with 500 μl of cell lysis buffer. To improve the quality of the immunoprecipitation experiment, first wash was last 30 minutes, at 4 °C on gentle shaking. Beads were finally resuspended with 20 μl 3X SDS sample buffer, and samples were loaded on 4–20% stain free SDS-PAGE pre-packed gel (Bio-RAD), for immunoblot analysis. Rabbit anti-HA and mouse anti-MYC (3724S and 2276, Cell Signaling) were used for detection of transfected HA-RIP2 or MYC-XIAP and corresponding mutants at 1:1000 dilution. Rabbit anti β-actin antibody at 1:1000 dilution was employed for normalization of total protein amount (8457, Cell Signaling). For revelation, secondary antibodies linked to fluorophores were used at 1:1000 dilution: goat anti-rabbit and goat anti-mouse linked to Alexa 488 and Alexa 647 respectively (A11008 and A32728, Thermo Fisher).

## Data availability

Coordinates for RIP2K-XIAP BIR2 complex are in the protein data bank) PDB with accession code 8AZA. The cryo-EM map has accession code EMD-15757. Movies have been deposited in EMPIAR with accession code ###. The NMR chemical shift assignments of XIAP BIR2_AG_^124-240^ have been deposited with the Biological Magnetic Resonance Bank (BMRB) under accession number 51600.

## Acknowledgements

We thank Katrin Rittinger (Francis Crick Institute, UK) for the kind plasmids gift. We acknowledge Alice Aubert and Martine Pelosse for support in using the Eukaryotic Expression Facility. We acknowledge Sarah Schneider and Wojtek Galej for support in using the EM facility at the EMBL in Grenoble. We thank the European Synchrotron Radiation Facility for provision of beam time on CM01. We acknowledge the EM platform in HD, for further data collections not here reported, but project connected. We thank the IT assistant of the IBS-EMBL shared processing cluster. This work used the platforms of the Grenoble Instruct-ERIC Center (ISBG; UAR 3518 CNRS-CEA-UGA-EMBL) within the Grenoble Partnership for Structural Biology (PSB), supported by FRISBI (ANR-10-INBS-0005-02) and GRAL, financed within the University Grenoble Alpes graduate school (Ecoles Universitaires de Recherche) CBH-EUR-GS (ANR-17-EURE-0003). Financial support from the IR-RMN-THC FR3050 CNRS is also acknowledged. We thank Caroline Mas for assistance and access to the biophysics platform. Molecular graphics and analyses were performed with UCSF ChimeraX, developed by the Resource for Biocomputing, Visualization, and Informatics at the University of California, San Francisco, with support from National Institutes of Health R01-GM129325 and the Office of Cyber Infrastructure and Computational Biology, National Institute of Allergy and Infectious Diseases.

## Author Contributions

E.P. and S.C. conceived and directed the project. M.L. and E.P performed cloning, biochemical and *in cellulo* experiments. E.P. took care of specimens preparation, screening and data collection at EM EMBL-Grenoble platform, M.H. performed data collection tilted mode at EM EMBL-Grenoble platform and both untilted and tilted mode at CM01, using in-house time. E.P. did data processing and structure determination. E.B.E. performed native mass spectrometry experiments. A.F. and B.B. performed NMR experiments. E.P. and S.C. wrote the manuscript with input from all authors.

