## Supporting Information for "Structural analysis shows that the BIR2 domain of E3 ligase XIAP binds across the RIP2 kinase dimer interface"

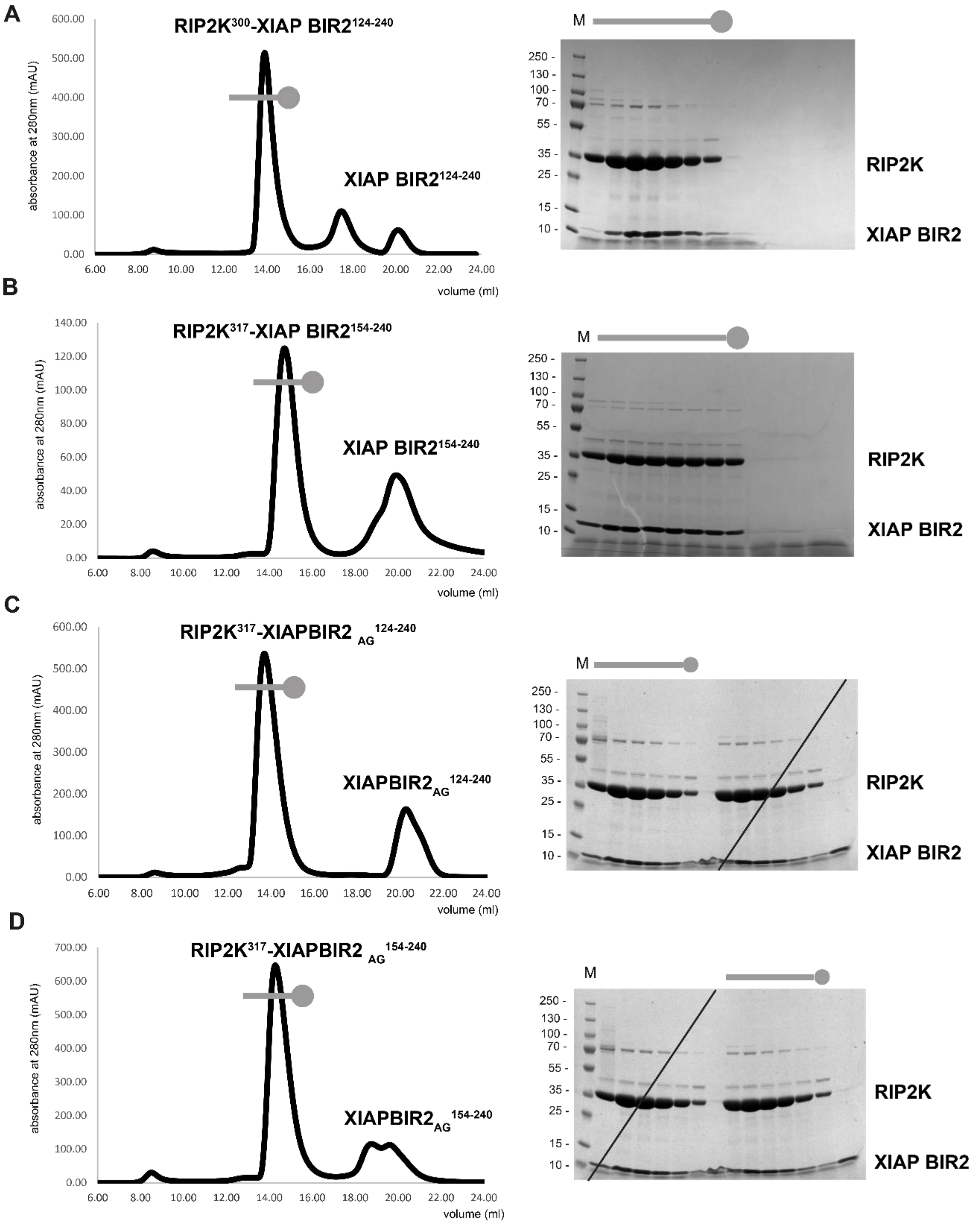


**Fig. S1. *In vitro* reconstitution of the RIP2K-XIAP BIR2 complex.** Size exclusion chromatography (SEC) profile of the 4 *in vitro* reconstituted RIP2K-XIAP BIR2 complexes togetherwith comassie-stained SDS-PAGE analysis of eluted fractions: (**A**) RIP2K$^{300}$-XIAP BIR2$^{124-240}$, (**B**) RIP2K$^{317}$-XIAP BIR2$^{154-240}$, (**C**) RIP2K$^{317}$-XIAP BIR2$_{AG}^{124-240}$, (**D**) RIP2K$^{317}$-XIAP BIR2$_{AG}^{154-240}$.The SEC profile of RIP2K$^{317}$-XIAP BIR2$^{154-240}$, already shown in Fig. 1B, it's here reported for completeness.

**A**

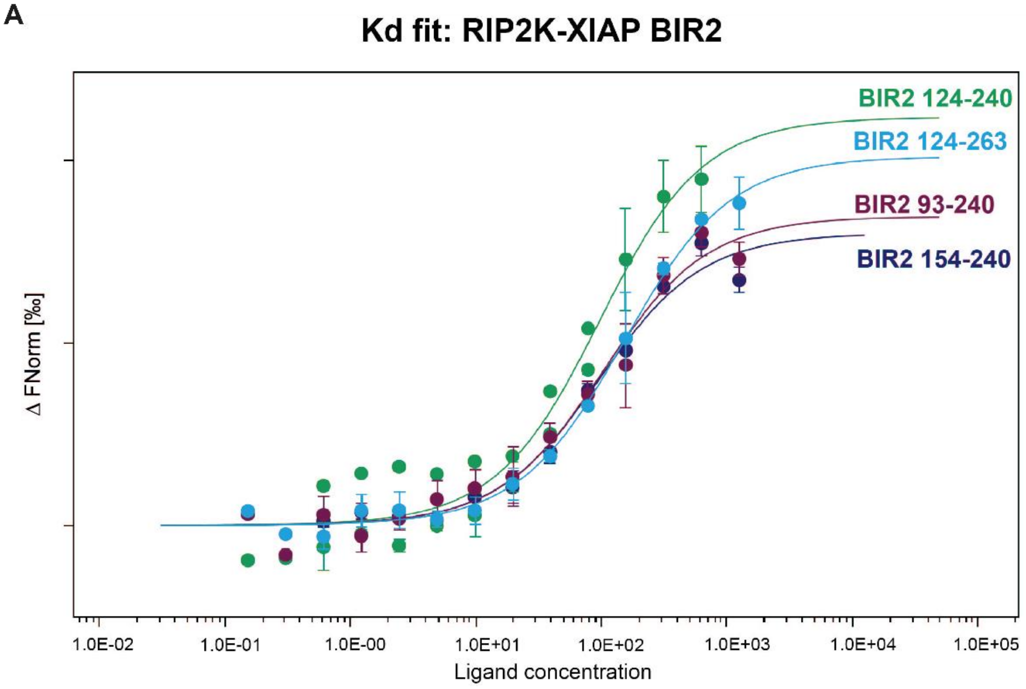


**B**

| Sample | Kd (nM) | Kd confidence |
|---|---|---|
| BIR2 154-240 | 85.6 | ±18.7 |
| BIR2 124-240 | 85.9 | ±24.9 |
| BIR2 93-240 | 92.9 | ±22.6 |
| BIR2 124-263 | 141.6 | ±15.4 |

**Fig. S2. RIP2K-XIAP BIR2 dissociation constants measured by MST.** (**A**) Dose-response curves of wt XIAP BIR2 constructs toward RIP2K. All experiments have been done in triplicates. Standard deviations are represented for each point in bars. (**B**) The table reports the Kd obtained for each sample. All the dose-response curves were fitted to a one-site binding model to obtain Kd value.

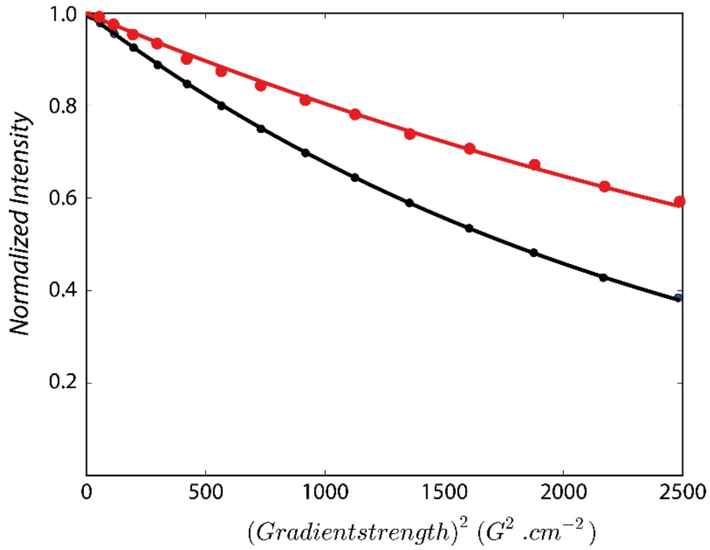


**Fig. S3. Translational diffusion properties of XIAP BIR2 (black) and the RIP2K$^{317}$-XIAP BIR2$_{AG}$$^{124\text{-}240}$ complex (red) measured by 1D $^1$H DOSY at 25°C and 850 MHz $^1$H frequency.** Exponential fitting of these decay curves results in diffusion coefficients of 13.6 $10^{-7}$ $m^2s^{-1}$ for XIAP BIR2 and 7.6 $10^{-7}$ $m^2s^{-1}$ for XIAP BIR2 : RIP2K. The translational diffusion coefficient provides a measure of the average particle size in solution. Upon interaction with RIP2K the apparent molecular size is increased by a factor of about 5.7, in good agreement with a 1:2 BIR2 : RIP2K stoichiometry.

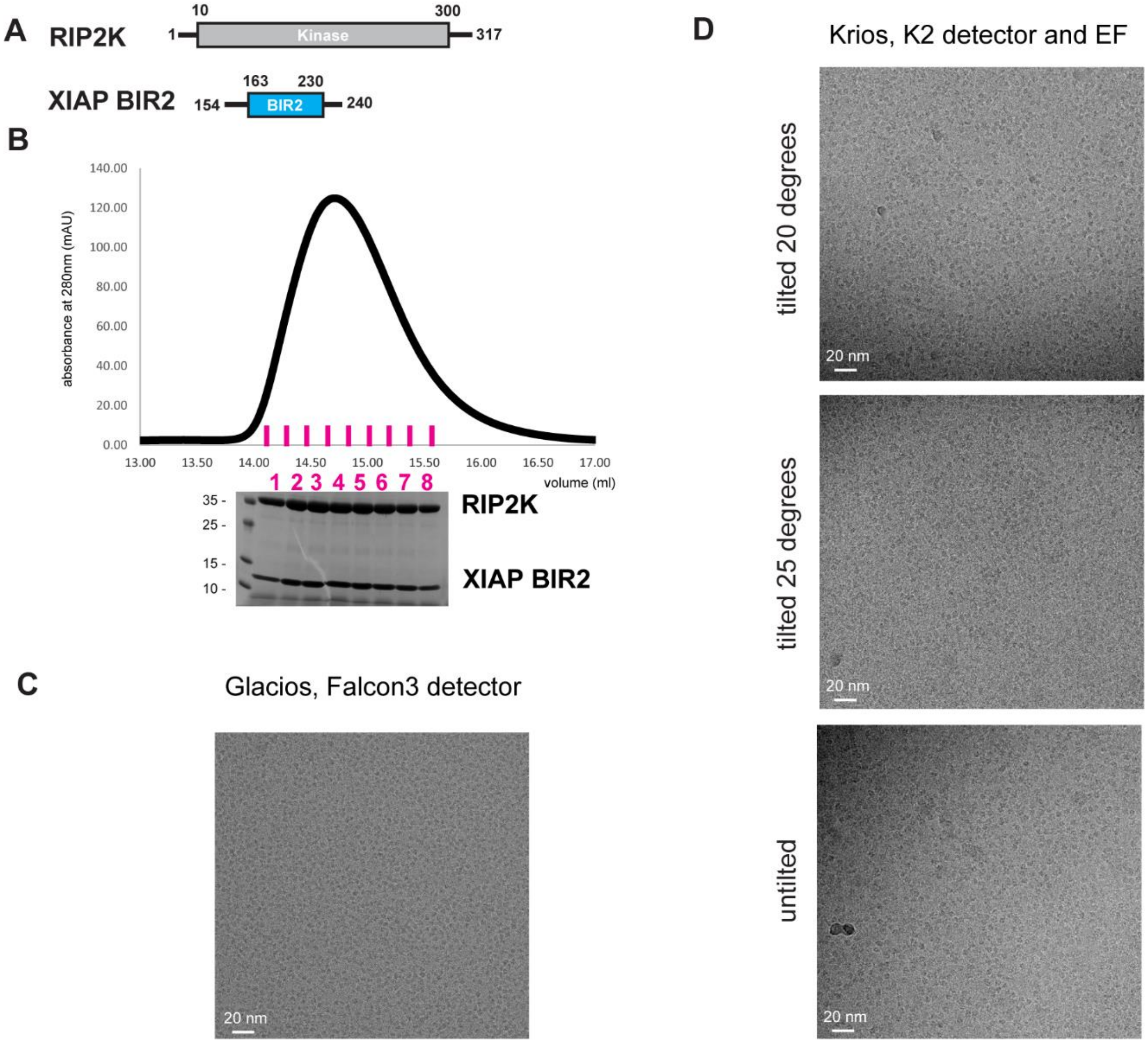


**Fig. S4. RIP2K$^{317}$-XIAP BIR2$_{AG}^{154\text{-}240}$ complex purification for cryo-EM.** (**A**) RIP2K and XIAP BIR2 constructs used for cryo-EM structure determination. (**B**) Zoom-in on the RIP2K$^{317}$-XIAP BIR2$_{AG}^{154\text{-}240}$ SEC profile (Fig.1SB); collected fractions are shown in magenta, fractions 3-6 have been used for cryo-EM specimen preparation. (**C**) Exemplary micrograph collected during specimen screening (Glacios TEM equipped with Falcon3, counting mode, -1 µm defocus, pixel size 0.94 px/Å, 40 e-/A2 total dose). (**D**) Exemplary micrographs collected at Krios in either tilted or untilted mode (EF=energy filter). Dose and pixel size are reported in the main text and in Table S2.

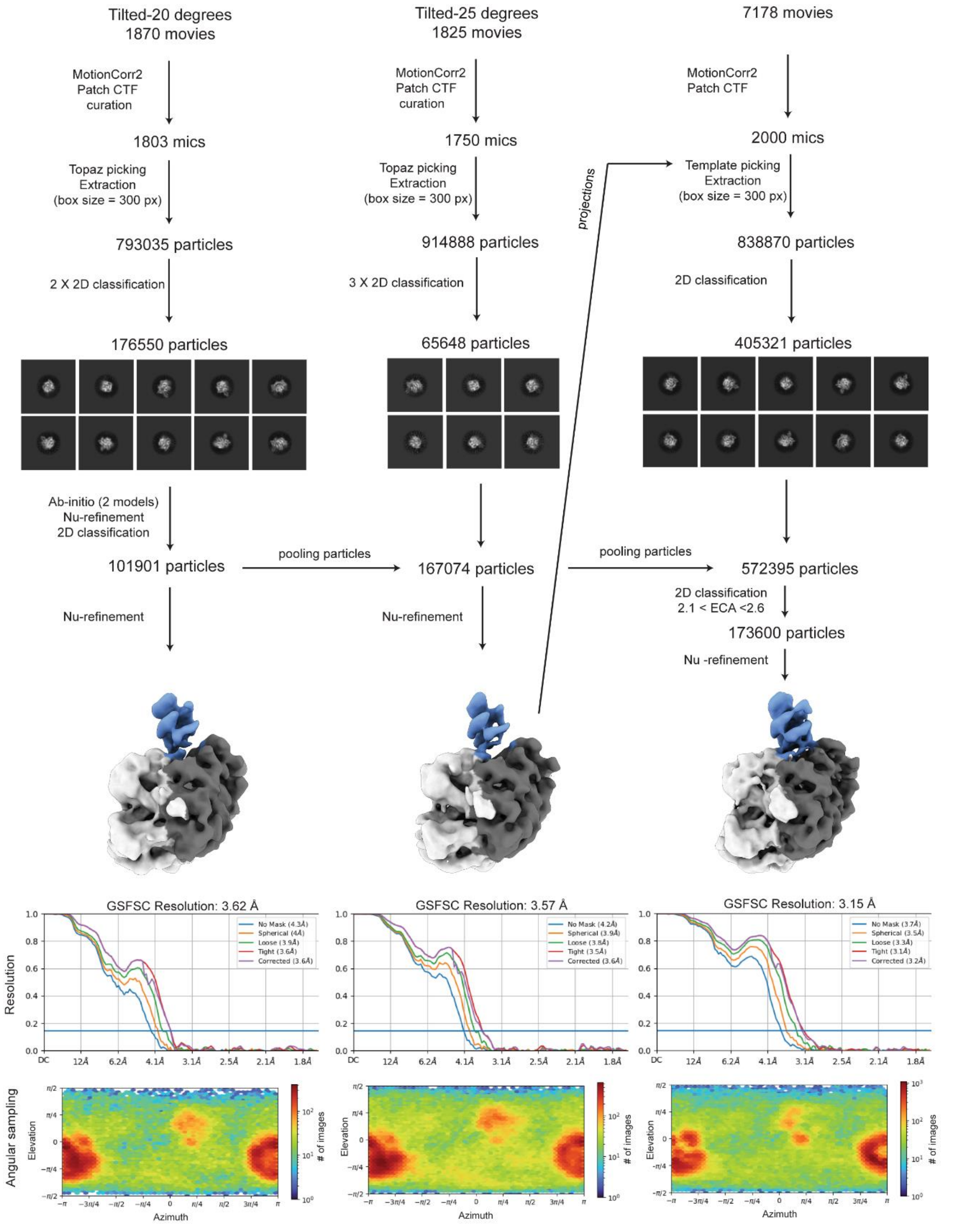


**Fig. S5. Cryo-EM processing workflow.** The scheme summarizes the cryo-EM processing to obtain the final map, as described in the Methods section. The unsharpened maps, the FSC threshold and the angular sampling schemes show how the map improved along the processing, without losing orientations and gaining in the density at the interaction between RIP2K (Kinase_A in light grey, Kinase_B in dark grey) and XIAP BIR2 (in light blue).

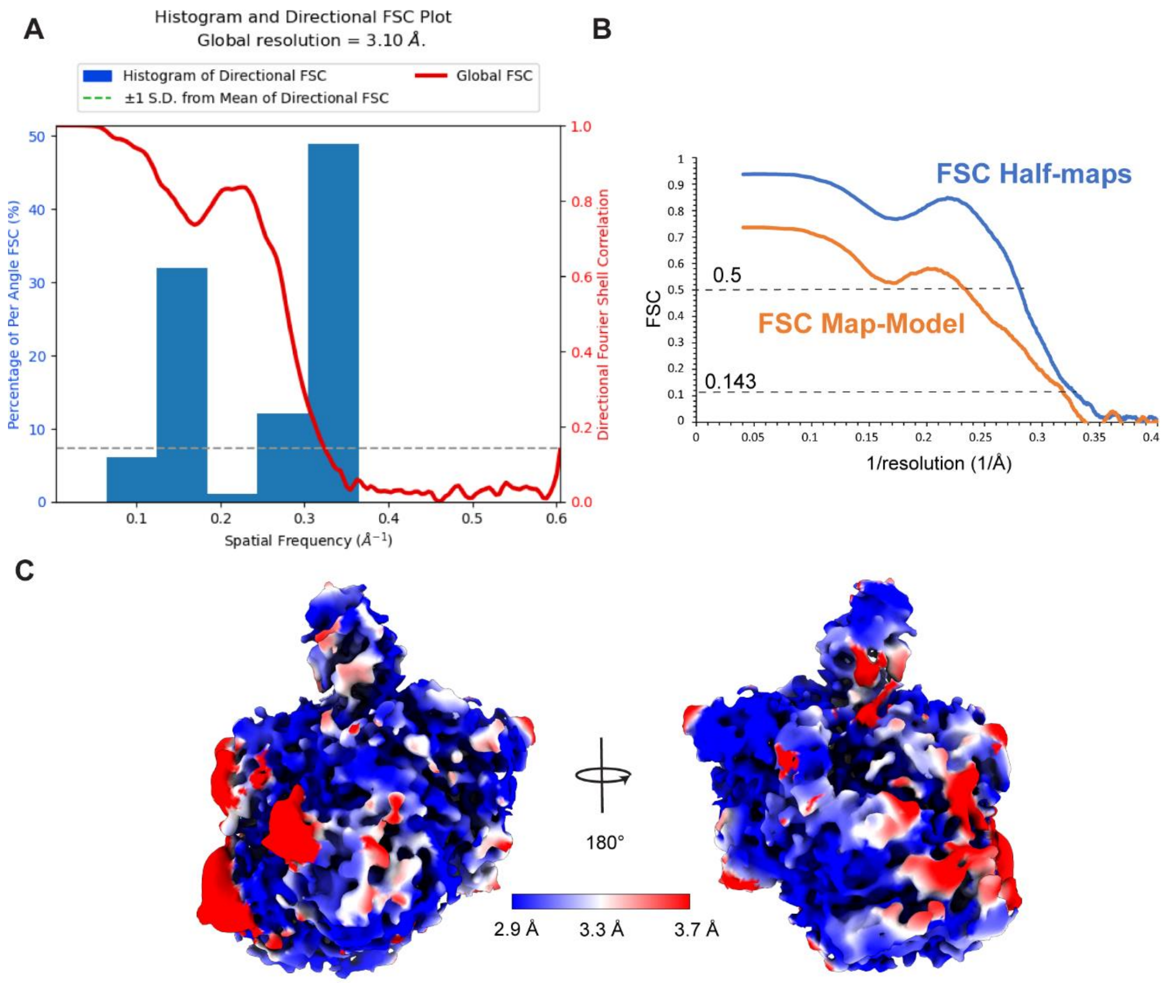


**Fig. S6. Local resolution of the RIP2K$^{317}$-XIAP BIR2$^{154\text{-}240}$ map.** (**A**) Directional FSC plot calculated in CryoSPARC. (**B**) Masked FSC half-maps and masked FSC map-model computed in Phenix (Afonine *et al*, 2018). (**C**) Cryo-EM density sharpened at B-factor -100 Å$^2$ of the RIP2K$^{317}$-XIAP BIR2$^{154\text{-}240}$ complex colored according to local resolution. Local resolution has been estimated in CryoSPARC.

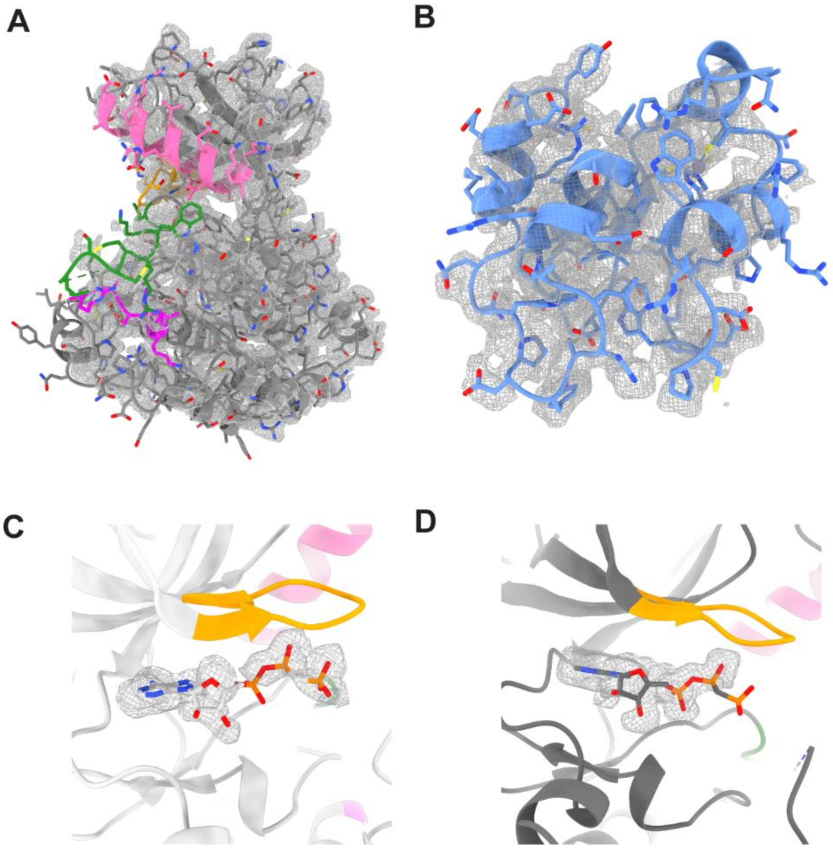


**Fig. S7. Fitting of RIP2K$^{317}$-XIAP BIR2$^{154\text{-}240}$ structure in the density map.** Fitting of (**A**) Kinase_A and (**B**) XIAP BIR2 domain in the cryo-EM density sharpened at B-factor -100 Å$^2$. (**C- D**) Cryo-EM density corresponding to the non-hydrolysable ATP analogue ACP in (**C**) Kinase_A and (**D**) Kinase_B. The fitting has been obtained by fitting in chimeraX the active structure of RIP2K (PDB ID: 5NG0)(Pellegrini *et al*, 2017). Chains and smaller elements are colored as in Fig. 3A.

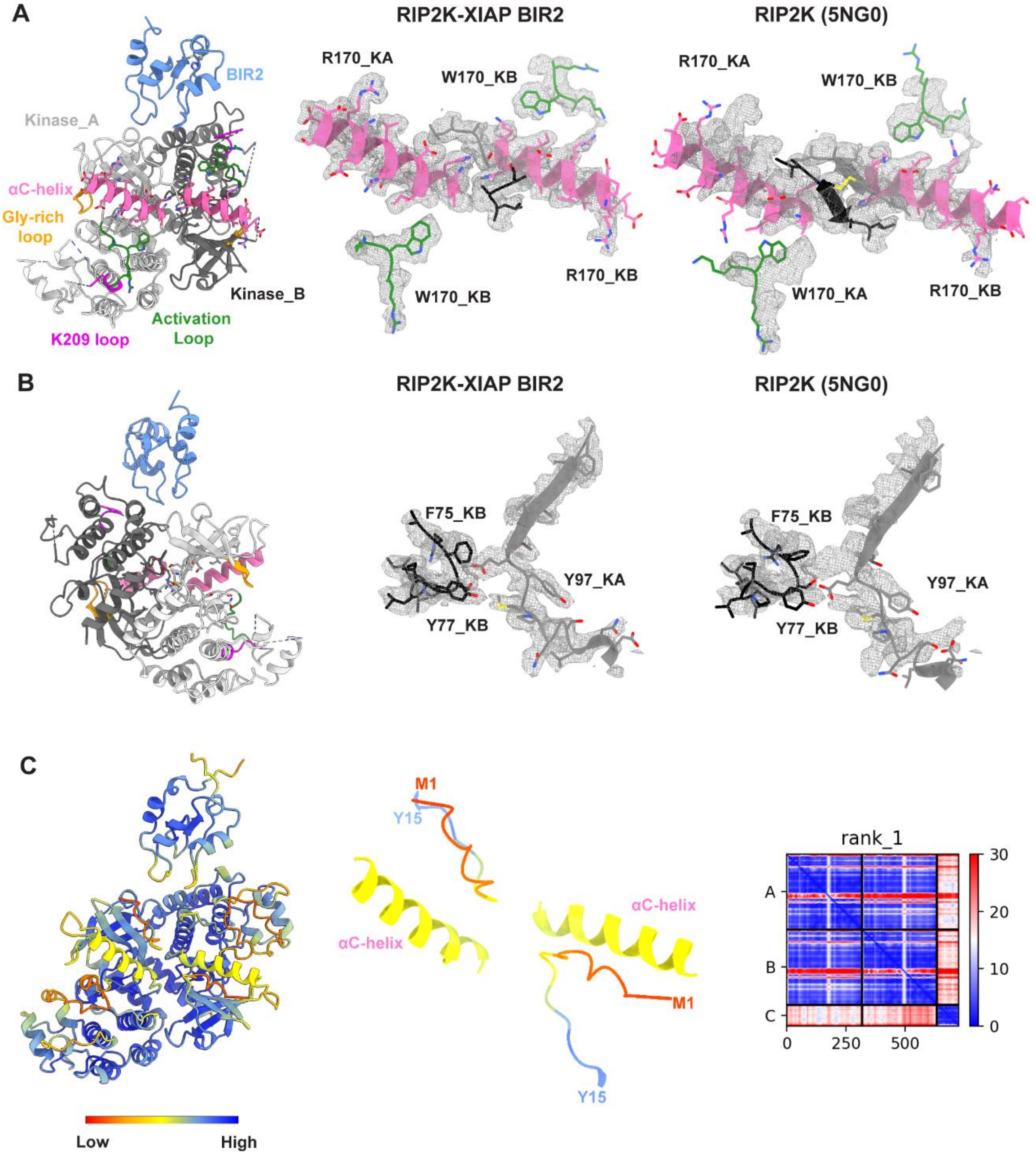


**Fig. S8. Flexible RIP2K N-terminus improves structure fitting in the EM density.** Comparison of fitting in the cryo-EM density between RIP2K dimer belonging to RIP2K$^{317}$-XIAP BIR2$^{154\text{-}240}$ structure (**A**) and active RIP2K structure (PDB ID: 5NG0) (Pellegrini *et al*, 2017)(**B**). Structures have been aligned on Kinase_B molecule of RIP2K$^{317}$-XIAP BIR2$^{154\text{-}240}$.

**(A) Left:** Ribbon representation of the RIP2K$^{317}$-XIAP BIR2$^{154\text{-}240}$ structure, with view on kinase αC-helixes and N-termini. Chains and smaller elements are colored as in Fig. 3A. **Middle**: Fitting of the αC-helixes (residues 57-72), the activation loops (residues 169-171) and kinase N-termini (residues 8-10) in the cryo-EM density. **Right**: Fitting in the density of the same protein elements belonging to the crystallographic RIP2K structure (PDB ID: 5NG0). In 5NG0, the N-terminus comprises residues 6-10. (**B**) **Left:** Ribbon representation of the RIP2K$^{317}$-XIAP BIR2$^{154-240}$ structure, rotated of 180 degrees. **Middle**: Fitting of residues 90-105 from Kinase_ A and 73-82 from Kinase_B in the cryo-EM density. **Right**: Fitting in the density of the same protein elements belonging to the crystallographic RIP2K structure (PDB ID: 5NG0). (**C**) **Left**: alphafold2 prediction of the RIP2K$^{317}$-XIAP BIR2$^{154-240}$ structure, colored according to prediction confidence (blue=100, yellow=70, orange=50, red=0). **Middle**: zoom-in on the αC-helixes and N-termini. **Right**: Predicted Aligned Error diagram (PAE) of alphafold2 prediction.

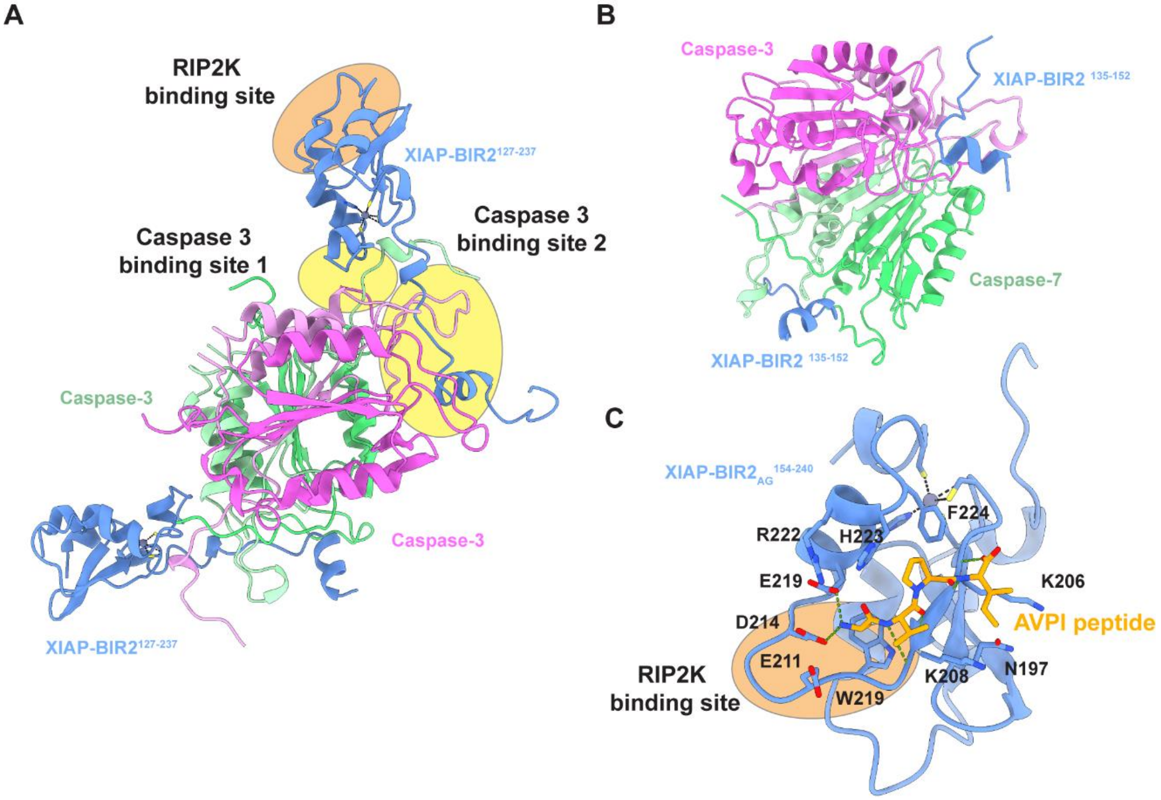


**Fig. S9. Binding of XIAP BIR2 domain to caspase 3, caspase 7 and the AVPI peptide. (A)** Ribbon representation of the XIAP BIR2-caspase 3 complex (PDB ID: 1i3o) (Riedl *et al*, 2001) The BIR2 domain of XIAP (colored in blue) binds backward to the catalytic subunit of caspase 3 compared to RIP2K binding (see the orange and yellow ovals). Using BIR2 residues N226, F228, R233 (Caspase 3 binding site 1) and the linker region (Caspase 3 binding site 2), XIAP interacts with both catalytic subunits of caspase 3 (colored in pink and green). Specifically, each XIAP BIR2 molecule (residues 127-237) interacts with both the large and small subunits of one caspase molecule (colored in dark and light pink respectively) and with a short stretch of the adjacent caspase molecule (colored in dark and light green). (**B**) Ribbon representation of the XIAP BIR2-caspase 7 complex (PDB ID: 1i51) (Chai *et al*, 2001). XIAP BIR2 interacts with both large and small domains of the catalytic subunit of caspase 7 (colored in dark pink/green and light pink/green respectively) using the linker region (residues 135-152, colored in blue). (**C**) Ribbon representation of XIAP BIR2 domain bound to AVPI peptide (PDB ID: 4j46)(Lukacs *et al*, 2013). The XIAP BIR2 residues involved in the binding to the peptide are highlighted in stick (IBM grove). As shown by the orange oval, the binding to RIP2K overlaps with the binding to the SMAC peptide.
Zn is represented in grey sphere. Metal ion bonds are represented in black dashed line, whilst H-bond are represented in green dashed line.

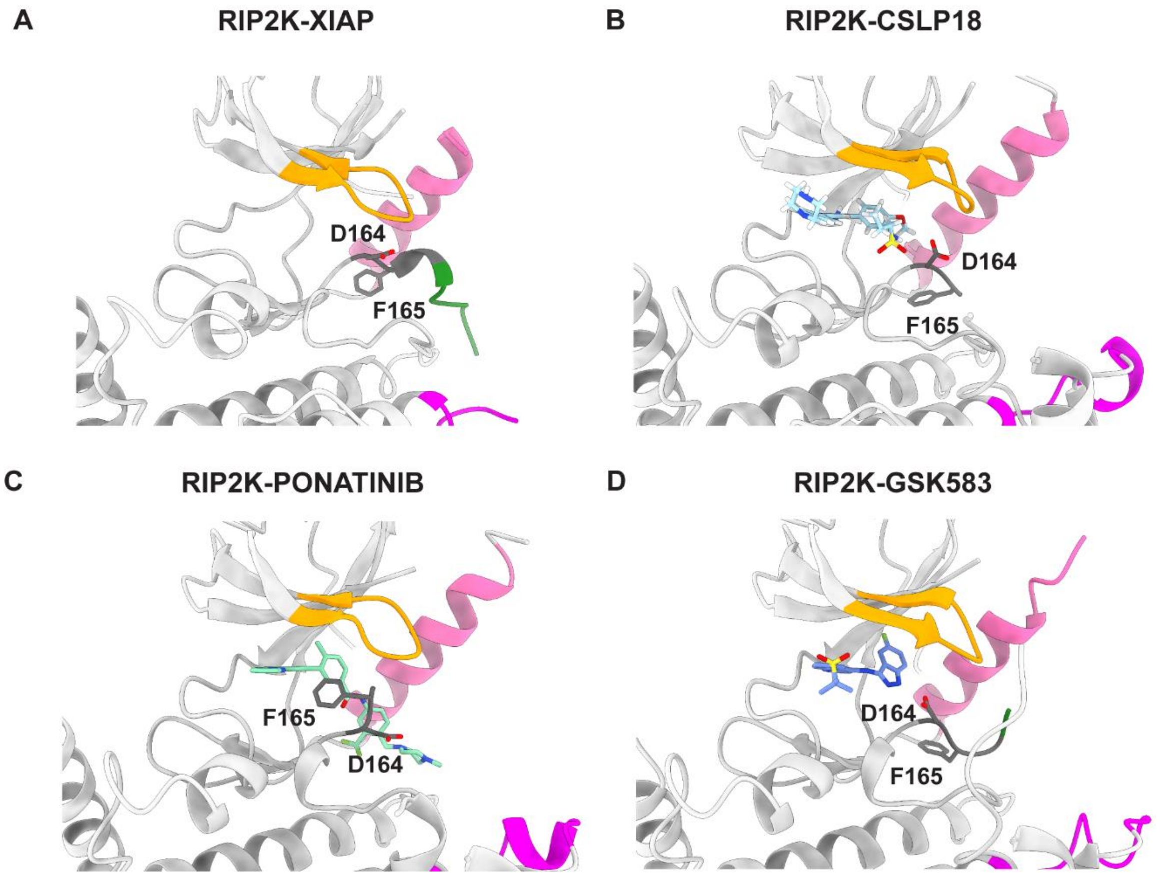


**Fig. S10. Conformation of DFG motif and activation loop in ligand bound RIP2K structures.** Ribbon representation of (**A**) RIP2K$^{317}$-XIAP BIR2$^{154\text{-}240}$ structure (this study) and RIP2K in complex with (**B**) CSLP18 (PDB ID: 6FU5) (Hrdinka *et al*, 2018), (**C**), Ponatinib (PDB ID: 4c8b) (Canning *et al*, 2015), (**D**) GSK583 (PDB ID: 5j7b) (Haile *et al*, 2016). Each figure shows the conformation of the DFG motif (residues 164-166, in stick) and the activation loop if present. The αC-helix is colored in pink, the Gly-rich loop in yellow, the activation loop in green and the K209 loop in magenta.

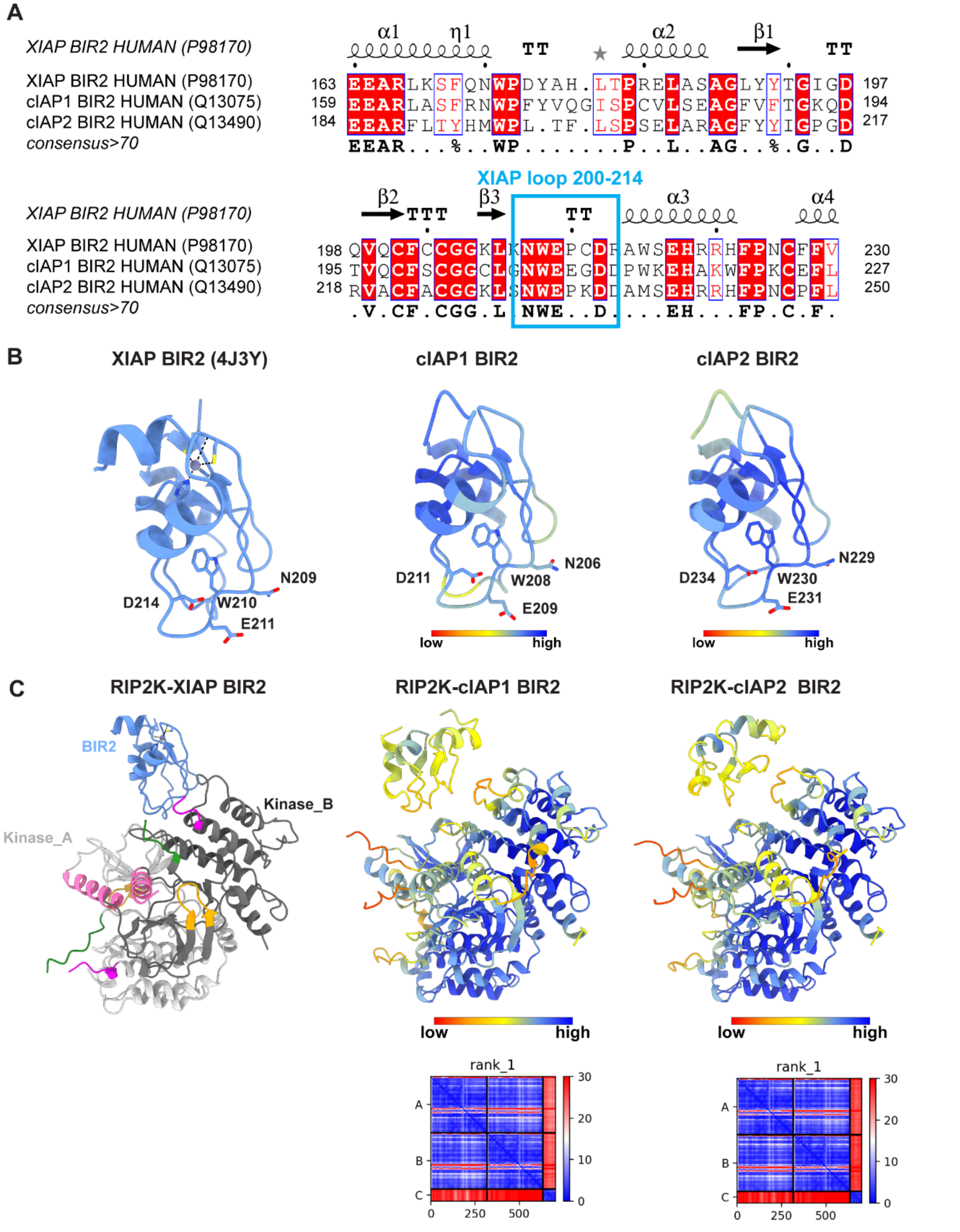


**Fig. S11. Alphafold2 predictions of cIAP1 BIR2, cIAP2 BIR2, RIP2K-cIAP1 BIR2 and RIP2K-cIAP2 BIR2 structures.** (**A**) Structure-based alignment of cIAPs BIR2 domains, within assigned Uniprot code. The alignment was computed using Clustal Omega (Madeira *et al*, 2022) and the output was used as input for ESPript 3.0, to generate the figure (Robert & Gouet, 2014). The secondary structure assignment of the XIAP BIR2 crystallographic structure is shown on top (PDB ID: 4J3Y)(Lukacs *et al*, 2013). Residues with consensus >70 are highlighted in red. The high similarity with the XIAP loop 200-214 is highlighted in blue (NWExxDx motif). (**B**) Ribbon representation of the crystallographic XIAP BIR2 structure and the alphafold2 predicted structured of cIAP1 BIR2 and cIAP2 BIR2 domains (from alphafold2 database). Side chains of the NWExxDx motif are shown. (**C**) Ribbon representation of the RIP2K$^{317}$-XIAP BIR2$^{154\text{-}240}$ structure (this paper) and the alphafold2 predicted structures of RIP2K-cIAP1 BIR2 and RIP2K-cIAP2 BIR2 complexes. Alphafold2 predicted models are coloured according to prediction confidence (blue=100, yellow=70, orange=50, red=0). The Predicted Aligned Error diagram (PAE) is shown for the structures computed by us.

**Table S1. Summary of Native MS results**

| Sample | Main oligomeric state | Experimental masses (Da) | Other oligomeric states | Experimental masses (Da) |
|---|---|---|---|---|
| **RIP2K$^{317}$-XIAP BIR2$^{154-240}$** | 2:1 | 84093 | 2:0, 2:2 | 73668 (Rip2K alone), 94518 |
| **RIP2K$^{300}$-XIAP BIR2$_{AG}^{154-240}$** | 2:2 | 90708 | 2:0, 2:1 | 70010 (Rip2K alone), 80359 |
| **RIP2K$^{317}$-XIAP BIR2$^{124-240}$** | 2:1 | 87461 | 2:0, 2:2, 2:3 | 73668 (Rip2k alone), 101254, 115047 |
| **RIP2K$^{300}$-XIAP BIR2$_{AG}^{124-240}$** | 2:2 | 97441 | none | - |

**Table S2. Cryo-EM data collection, refinement and validation statistics**

| | RIP2K-XIAP BIR2 (PDB-8AZA, EMD-15757) |
|---|---|
| **Data collection and processing** | |
| Magnification | 165,000 x |
| Voltage (kV) | 300 |
| Electron exposure (e–/Å$^2$) | 47.7 and 45.7 for tilted dataset |
| Defocus range (μm) | -0.8 to -2.5 |
| Pixel size (Å) | 0.87 |
| Symmetry imposed | - |
| Initial particle images (no.) | 2,546,793 |
| Final particle images (no.) | 173,600 |
| Map resolution (Å) | 3.15 |
| FSC threshold | 0.143 |
| Map resolution range (Å) | 2.9 -5.488 (25$^{th}$ to 75$^{th}$ percentile) |
| **Refinement and Validation** | |
| Initial model used (PDB code) | 5NG0, 4C8B and 4J3Y |
| Map sharpening *B* factor (Å$^2$) | -100 |
| Model composition | |
| Non-hydrogen atom | 5031 |
| Protein residues | 614 |
| Waters | 0 |
| Ligands | 1 (Zn) |
| *B* factor (Å$^2$) | 165 (masked) |
| R.m.s. deviations | |
| Bond lengths (Å) | 0.004 |
| Bond angles (°) | 0.758 |
| Validation | |
| MolProbity score | 2.26 |
| Clashscore | 26.56 |
| Poor rotamers (%) | 0.77 |
| Ramachandran plot | |
| Favored (%) | 95.07 |
| Allowed (%) | 4.93 |
| Disallowed (%) | 0 |
| Model vs Data | |
| CC (mask) | 0.67 |
| CC (box) | 0.83 |
| CC (peaks) | 0.58 |
| CC (volume) | 0.66 |
| Mean CC for ligands | 0.50 |

**Table S3. Oligos for cloning and mutagenesis**

**Oligos used for XIAP extensions:**

| Constructs | Plasmid | Oligo |
|---|---|---|
| 124-240 Fw | pET49 XIAP 154-240 | 5'-GCAGACTATCTTTTGAGAACTGGGCAGGTTGTAGATATATCAGACACCATATACCCGAGGAACCCTGCCATG-3' |
| 124-240 Rev | pET49 XIAP 154-240 | 5'-CAAAAGATAGTCTGCATGTGTCTCAGATGGCCTGTCTAAGGCAAAATGATCTCTGGGTCCCTGAAAGAGGACTTC-3' |
| 93-240 Fw | pET49 XIAP 124-240 | 5'-ACAAATTCTGGTATCCAGAATGGTCAGTACAAAGTTGAAAACTATCTGGGAAGCAGAGATCATTTTGCCTTAGACAG-3' |
| 93-240 Rev | pET49 XIAP 124-240 | 5'-GATACCAGAATTTGTAGACTGCGTGGCACTATTTTCAAGATAAAAGCCGTTGATGGGTCCCTGAAAGAGGAC-3' |
| 124-263 Fw | pET49 XIAP 124-240 | 5'-TTTCCCAAATTCAACAAATCTTCCAAGAAATCCATCCATGGCATAACGCTCTGGTGCCACG-3' |
| 124-263 Rev | pET49 XIAP 124-240 | 5'-GTTGAATTTGGGAAATTCCTATCAGAACTCACAGCATCAGATTCACTTCGAATATTAAGATTCCG-3' |

**Oligos used for RIP2K extension (on pFastBac-HTB RIP2):**

| Construct | Oligo |
|---|---|
| 0-317 Fw | 5'-GTTATTCAGCTAAAGAAAACAAAGTTACAGAGTTGACTCGAGAAGCTTGTCGAG-3' |
| 0-317 Rev | 5'-CTTTAGCTGAATAACAGCTTCAAGAAAAGTTATCTCTTCAAATGTTCTCAAAACTG-3' |

**Oligos used for Myc-3c hXIAP cloning into pCDNA3:**

| Construct | Oligo |
|---|---|
| Fw | 5'-TAGCTGGATCCATGGAACAAAAACTCATCTCAGAAGAG<br>GATCTGCTGGAAGTTCTGTTCCAGGGGCCCGCTAGTGCTA<br>GTGCTAGTATGACTTTTAACAGTTTTGAAGGATC-3' |
| Rev | 5'-TATCGACTCGAGTTATTAAGACATAAAAATTTTTTGCTTGAAAG-3' |

**Oligos for XIAP mutagenesis (on pCDNA3-Myc-3c hXIAP):**

| Construct | Oligo |
|---|---|
| XIAP Y176A | 5'-CAGAACTGGCCAGACGCTGCTCACCTAACCCC-3' |
| XIAP H178A | 5'-CCAGACTATGCTGCCCTAACCCCAAGAG-3' |
| XIAP N209A | 5'-GTGGTGGAAAACTGAAAGCTTGGGAACCTTGTGATC-3' |
| XIAP E211A | 5'-CTGAAAAATTGGGCACCTTGTGATCGTGCCTGG-3' |
| XIAP C213A | 5'-GAAAAATTGGGAACCTGCTGATCGTGCCTGGTC-3' |
| XIAP D214A | 5'-GGGAACCTTGTGCTCGTGCCTGGTCAG-3' |
| XIAP R215A | 5'-GGGAACCTTGTGATGCTGCCTGGTCAGAACAC-3' |

**Oligos for RIP2 mutagenesis (on pCDNA3-HA-RIP2):**

| Construct | Oligo |
|---|---|
| RIP2-R36L | 5'-GTCGTCCGCCCTCCACGCAGACTG-3' |
| RIP2-D39L | 5'-CCGCCACGCACTCTGGCGCGTCCAG-3' |
| RIP2-R41L | 5'-CGCAGACTGGCTCGTCCAGGTGGC-3' |
| RIP2-R39L/R41L | 5'-CGCAGACTGGCTCGTCCAGGTGGC-3' (ON RIP2-R49L) |
| RIP2-N137L | 5'-GGTGTAAATTACCTGCACCTTATGACTCCTCCTTTAC-3' |
| RIP2-N137L/N133L | 5'-GCCCTTGGTGTACTTTACCTGCACCTTATGACTCCTCCTTTACTTC-3' |
| RIP2-K209A | 5'-CAAGGGCCAGTATCGCGCACGATATATATAGCTATG-3' |
| RIP2-K209R | 5'-CAAGGGCCAGTATCCGGCACGATATATATAGCTATG-3' |
| RIP2-E279L | 5'-GGGCACAAAATCCAGATCTAAGACCATCTTTCTTAAAATG-3' |
| RIP2-S282L | 5'-CACAAAATCCAGATGAAAGACCACTTTTCTTAAAATGTTTAATAGAAC-3' |
| RIP2-K285A | 5'-CAAGGGCCAGTATCGCGCACGATATATATAGCTATG-3' |
| RIP2-K285L | 5'-GATGAAAGACCATCTTTCTTACTATGTTTAATAGAACTTGAACC-3' |
| RIP2-W170A | 5'-GATTTTGGTTTATCAAAGGCGCGCATGATGTCCCTCTC-3' |
| RIP2-R171A | 5'-GGTTTATCAAAGTGGGCCATGATGTCCCTCTCACAG-3' |
